# hnRNPM and ELAVL1 control type I interferon induction by promoting IRF3 phosphorylation downstream of both cGAS and RIG-I

**DOI:** 10.1101/2023.06.23.545108

**Authors:** Alexander Kirchhoff, Anna-Maria Herzner, Christian Urban, Antonio Piras, Robert Düster, Julia Wegner, Agathe Grünewald, Thais M. Schlee-Guimarães, Katrin Ciupka, Petro Leka, Robert J. Bootz, Ann Kristin de Regt, Beate M. Kümmerer, Maria Hønholt Christensen, Florian I. Schmidt, Claudia Günther, Hiroki Kato, Eva Bartok, Gunther Hartmann, Matthias Geyer, Andreas Pichlmair, Martin Schlee

**Author notes:** To whom correspondence should be addressed., (M.S.). Department of Cancer Immunology and Immune Modulation, Boehringer Ingelheim Pharma GmbH & Co. KG, Biberach an der Riß, Germany.

## Abstract

RIG-I and cGAS are crucial sensors of viral nucleic acids and induce type I IFNs via TBK1/IKK and IRF3. Here, we have identified hnRNPM as a novel positive regulator of IRF3 phosphorylation and type I IFN induction downstream of both cGAS and RIG-I. Combining interactome analysis and genome editing, we further identified ELAVL1 as an immune-relevant interactor of hnRNPM. Depletion of hnRNPM or ELAVL1 impaired type I IFN induction by HSV-1 and SeV. In addition, we found that hnRNPM and ELAVL1 interact with TBK1 and NF-kB p65. Confocal microscopy revealed cytosolic and perinuclear interactions between hnRNPM, ELAVL1, and TBK1. To our knowledge, hnRNPM and ELAVL1 represent the first non-redundant signaling components merging the cGAS-STING and RIG-I-MAVS pathways, thus representing a novel platform that fuels antiviral defense.

## INTRODUCTION

Recognition of non-self nucleic acids is an essential defense mechanism of the innate immune system ^1^. As multiple pathogens exploit the cytosol as a replication niche, it is constantly monitored for the presence of pathogen-associated molecular patterns (PAMPs) by specialized pattern recognition receptors (PRRs). In the cytosol, cyclic GMP-AMP (cGAMP) synthase (cGAS) constitutes the principal type I interferon (IFN)-inducing receptor of double-stranded DNA (dsDNA), whereas retinoic acid inducible gene I (RIG-I) and melanoma differentiation associated gene 5 (MDA5) are the predominant sensors of cytosolic dsRNA species ^2, 3^. All three receptors signal via the formation of multimeric protein complexes leading to interferon regulatory factor 3 (IRF3)-dependent type I IFN induction. Detection of cytoplasmic dsDNA activates cGAS to produce 2’3’-cGAMP, a second messenger that binds to the endoplasmic reticulum (ER)-resident adaptor protein stimulator of interferon genes (STING) ^4, 5^. Upon activation, STING multimerizes and translocates from the ER to the Golgi compartment. STING clustering provides a suitable surface for the recruitment of multiple TANK-binding kinase 1 (TBK1) molecules ^6–8^. Local accumulation of TBK1 molecules induces *trans*-autophosphorylation in the activation loop at Ser172, followed by TBK1-dependent phosphorylation of STING at Ser366 in the conserved pLxIS motif (p, hydrophilic residue; x, any residue) ^9, 10^. Binding of tri-/di-phosphorylated dsRNA and long dsRNA species by RIG-I and MDA5, respectively, induces caspase recruitment domain (CARD)-driven multimerization and interaction with mitochondrial antiviral-signaling protein (MAVS) at the cytoplasmic portion of the outer mitochondrial membrane, where MAVS is phosphorylated in the pLxIS motif at Ser442 by TBK1, inhibitor of kappa B kinase-χ (IKKχ), or IKKβ ^10–14^. Thus, the signaling pathways downstream of cGAS and RIG-I/MDA5 converge at the stage of TBK1/IKK. Following, IRF3 is recruited to the phosphorylated pLxIS motifs of STING or MAVS, leading to homodimerization and nuclear translocation of dimeric IRF3 ^10, 15, 16^. Additionally, STING and MAVS induce the activation of nuclear factor kappa-light-chain-enhancer of activated B cells (NF-κB) via IKKα/IKKý ^17^. In concert with other transcription factors, IRF3 and NF-κB drive the expression of type I IFNs and pro-inflammatory cytokines.

Similar to cGAS and RIG-I/MDA5, activation of Toll-like receptor (TLR) 3 or TLR4 leads to the TBK1-dependent phosphorylation of an adaptor protein (TIR-domain-containing adapter-inducing IFN-β (TRIF)) at a consensus pLxIS motif, which serves as a binding site for IRF3 and is thus essential for type I IFN induction. By contrast, stimulation of myeloid differentiation primary response 88 (MyD88)-dependent TLRs such as TLR1/2 also induces TBK1 phosphorylation but does not lead to an IRF3-dependent type I IFN response ^18, 19^. Phosphorylation of TBK1 is therefore necessary but not sufficient to activate IRF3, as this further requires the engagement of appropriate adaptor proteins. Although ligand preferences and molecular functions of cGAS and RIG-I have now been described in detail ^20^, it is incompletely understood how signaling proteins shared by unrelated pathways integrate different input signals and induce PRR-specific gene expression programs. Accumulating evidence suggests that central kinases such as the canonical (IKKα, IKKý) and non-canonical IKKs (TBK1, IKKϵ) are targeted to distinct signaling complexes in a PRR-dependent manner, thereby enabling the cell to specify and spatiotemporally regulate the antiviral response.

Heterogeneous nuclear ribonucleoprotein M (hnRNPM) has been predominantly described in the context of pre-mRNA splicing, cancer biology, or muscle differentiation ^21–24^. In addition, hnRNPM was described to be targeted by the 3C proteases of Coxsackievirus B3 (CVB3) and Poliovirus (PV) leading to an increased replication of these viruses ^25^ and reported to suppress the expression of a cluster of immune-related genes in RAW 264.7 cells ^26^. Curiously, hnRNPM was shown to restrict growth of *Listeria monocytogenes* (*L. monocytogenes*) and certain alphaviruses (Semliki Forest virus (SFV), Chikungunya virus (CHIKV)) ^27, 28^. Recently, interactions between hnRNPM and ORF3B, a potent type I IFN antagonist encoded by SARS-CoV1, have been identified in high-throughput screens ^29, 30^. However, the molecular role of hnRNPM in innate antiviral immunity remains elusive.

In this study, we found that hnRNPM promotes the phosphorylation of IRF3 and expression of type I IFNs induced both cGAS and RIG-I in a non-redundant manner. Through a combined affinity purification followed by mass spectrometry (AP-MS)-based RNAi screening approach, we identified ELAV-like protein 1 (ELAVL1; also known as HuR) as an interactor of hnRNPM crucial for IRF3 phosphorylation and subsequent induction of type I IFNs and activation of NF-κB. Moreover, we provide evidence that hnRNPM, ELAVL1, TBK1, and NF-κB p65 form a multiprotein complex that fuels type I IFN induction by linking cGAS and RIG-I signaling at the stage of IRF3 activation.

## RESULTS

### hnRNPM promotes both cGAS-STING- and RIG-I-dependent production of type I IFNs

The 78 kDa hnRNPM is a multidomain RNA-binding protein with binding preferences towards G/U-rich intron mRNA and consists of a non-classical bPY nuclear localization sequence, three RNA-recognition motifs (RRMs), a glycine/methionine-rich region (GMG), and a methionine/arginine repeat motif (MR) ^22^. As hnRNPM is cleaved by certain viral proteases and since viruses often attempt to inactivate components of the nucleic acid-sensing pathways to evade immune surveillance, we set out to explore the role of hnRNPM in host defense. First, we analyzed degradation of hnRNPM by CVB3 and PV 3C proteases. As expected, co-expression of FLAG-tagged variants of these 3C proteases with GFP-tagged hnRNPM in HEK293FT cells potently reduced hnRNPM expression (Figure 1A). To elucidate whether hnRNPM is involved in nucleic acid-sensing, we used a human monocytic cell line (THP1) with an integrated IFN-stimulated response element (ISRE) reporter as a model system. Usage of a well-characterized ISRE reporter largely avoids detection of post-transcriptional regulation (splicing, UTR-dependent degradation) and therefore mirrors the activation of IRF transcription factors. Similar to other research teams, we did not obtain homozygous knockout (KO) clones after depletion of *HNRNPM* by CRISPR-Cas9 in THP-1, suggesting that full depletion of hnRNPM is not tolerated by the cell ^26^. Thus, we generated THP-1 hnRNPM knockdown (KD) cells. A moderate KD of hnRNPM was confirmed by qPCR and did not impair cellular viability (Figures 1B–1C). Of note, given only a moderate KD of hnRNPM, we observed an unexpectedly strong inhibition of the ISRE reporter in hnRNPM KD cells after stimulation with agonists for RIG-I (5’ppp-dsRNA), cGAS (plasmid DNA: pDNA, Y-DNA: G_3_-YSD), or STING (2’3’-cGAMP) (Figure 1D). Here, pathway inhibition was proportional to reduction of hnRNPM expression, while ISRE reporter activity induced by stimulation of TLR8 (TL8-506) or IFNα receptor (IFNAR) complex was not impaired by KD of hnRNPM (Figures 1D–1E). Next, we analyzed the role of hnRNPM after infection with *L. monocytogenes,* herpes simplex virus type 1 (HSV-1), and Sendai virus (SeV), which are known to be sensed by cGAS, STING, and/or RIG-I ^31–34^. Compellingly, hnRNPM KD significantly reduced ISRE reporter induction during infection with *Listeria,* HSV-1, or SeV (Figures 1F–1H). To further investigate the antimicrobial activity of hnRNPM, we asked whether hnRNPM regulates a signaling step downstream of cGAS and RIG-I. We thus analyzed phosphorylation of IRF3 at Ser396 (pIRF3-Ser396) induced by cGAS or RIG-I stimulation. Intriguingly, pIRF3-Ser396 levels elicited by cGAS or RIG-I were markedly reduced in THP-1 hnRNPM KD cells (Figure 1I). By contrast, the IFNα-induced phosphorylation of signal transducer and activator of transcription 1 (STAT1) at Tyr701 (pSTAT1-Tyr701) occurred independently of hnRNPM (Figure S1A), further indicating that hnRNPM functions upstream of the IFNAR complex in the ISG induction cascade. In summary, these data suggest that hnRNPM controls both the cGAS- and RIG-I-mediated induction of type I IFNs by promoting IRF3 activation.

**Figure 1:**
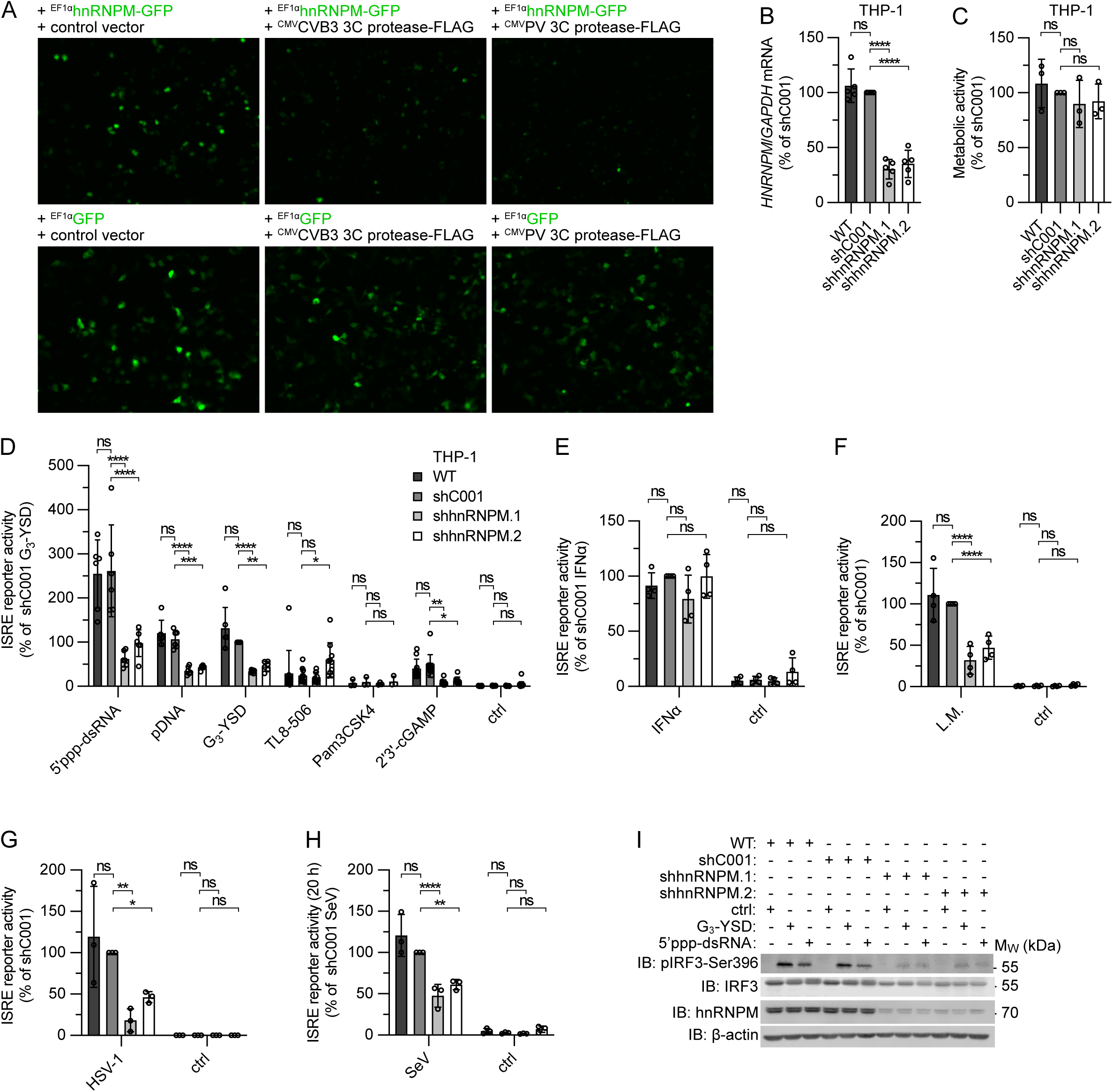
hnRNPM promotes type I IFN induction downstream of both cGAS and RIG-I. (A) Fluorescence microscopy of HEK293FT cells transiently transfected with hnRNPM-GFP (top) or GFP (bottom) together with a control vector or CVB3 3C protease-FLAG- or PV 3C protease-FLAG-expressing vectors (CMV or EF1α promotor). (B) qPCR of *HNRNPM* in THP-1 WT and cells expressing control shRNA (shC001) or hnRNPM-specific shRNAs (shhnRNPM.1, shhnRNPM.2). (C) MTT assay of the cells depicted in (B). (D) ISRE reporter activation in the cells depicted in (B) 20 h after stimulation of RIG-I with 5’ppp-dsRNA (0.1 µg/ml), of cGAS with pDNA (0.1 µg/ml) or G_3_-YSD (0.5 µg/ml), of TLR8 with TL8-506 (1.0 µg/ml), of TLR1/2 with Pam3CSK4 (0.5 µg/ml), or of STING with 2’3’-cGAMP (10 µg/ml). ctrl, non-stimulated. (E) ISRE reporter activation in the cells depicted in (B) 20 h after stimulation of IFNAR with IFNα (1000 U/ml). ctrl, non-stimulated. (F) ISRE reporter activation in the cells depicted in (B) 24 h after infection with *L. monocytogenes* (L.M., MOI 1). ctrl, non-stimulated. (G) ISRE reporter activation in the cells depicted in (B) 24 h after infection with HSV-1 (MOI 5). ctrl, non-stimulated. (H) ISRE reporter activation in the cells depicted in (B) 24 h after infection with Sendai virus (SeV) (MOI 1). ctrl, non-stimulated. (I) Immunoblot analysis of the cells depicted in (B) 3 h after stimulation with of cGAS with G_3_-YSD (0.5 µg/ml) or RIG-I with 5’ppp-dsRNA (0.1 µg/ml). ctrl, non-stimulated. One representative experiment of two independent experiments is shown. For (B)–(C): mean ± SD, one-way ANOVA, Dunnett’s multiple comparisons test. For (D)–(H): mean ± SD, two-way ANOVA, Dunnett’s multiple comparisons test.

### hnRNPM interacts with known innate immunity-associated factors

To further analyze the role of hnRNPM in innate immunity and to identify functionally important interactors that promote signaling downstream of both cGAS-STING and RIG-I-MAVS, we performed AP-MS in combination with an RNAi screen (Figure 2A). To this aim, we generated THP-1 cells stably expressing C-terminally GFP-tagged hnRNPM or GFP (control) (Figure 2A). We isolated cell populations with moderate GFP expression levels using fluorescence-activated cell sorting (FACS) to avoid artifacts caused by overexpression. Of note, expression of the hnRNPM-GFP fusion was weaker compared to endogenous hnRNPM (Fig. S1B). Next, we affinity-purified GFP from lysates of non-stimulated cells expressing GFP or hnRNPM-GFP and identified bound proteins by LC-MS/MS (Figure 2A). RNA-binding proteins associated to the ribosome (cluster (C) 1) and RNA splicing machinery (C2) were the most abundantly enriched interactors of hnRNPM (Figures 2B and S2; Table S1). In addition, hnRNPM interacted with constituents of the protein folding machinery (C3), mitochondrial transport proteins (C4), and proteins involved in immune system-related processes (C5) or cytokine signaling (C8) (Figure S2). Interestingly, we also observed interactions between hnRNPM and proteins that have already been associated with positive regulation of type I IFN expression, including ATP-dependent RNA helicase A (DHX9), pre-mRNA-splicing factor ATP-dependent RNA helicase DHX15 (DHX15), ELAVL1, DNA-dependent protein kinase catalytic subunit (PRKDC; also known as DNA-PKcs), DDX3X, and zinc finger CCCH-type antiviral protein 1 (ZC3HAV1) (Figure 2B) ^35–42^. Notably, we also observed interactions of hnRNPM with IKKβ (IKBKB), which is known to promote type I IFN induction by phosphorylating MAVS in the pLxIS motif (Figure 2B) ^10^.

**Figure 2:**
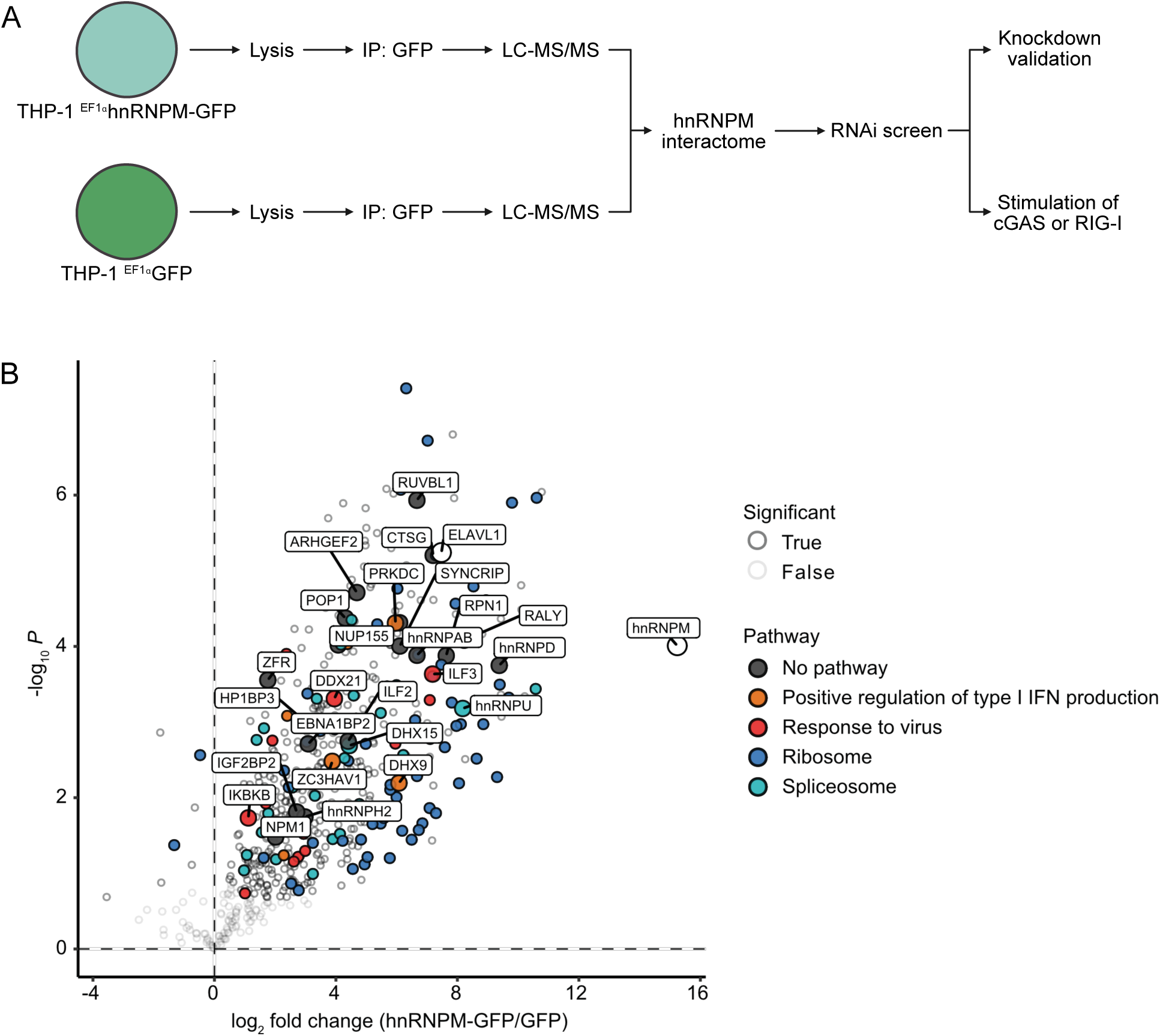
Interactome of hnRNPM. (A) Schematic workflow of the combined AP-MS-RNAi screening approach to identify immune-relevant interactors of hnRNPM. (B) hnRNPM-GFP and GFP were immunoprecipitated from lysates of non-stimulated THP-1 cells and bound proteins were identified by LC-MS/MS. The Volcano plot shows the log_2_ fold change enrichment of proteins detected in the hnRNPM-GFP vs. GFP affinity purification, plotted against the statistical significance (-log_10_ *P*). Significantly enriched proteins were determined by two-sided Welch’s t-tests (S0 = 0.1, permutation-based FDR < 0.05). Colors indicate pathway annotations and the majority of labeled proteins were functionally evaluated by RNAi.

### ELAVL1 is a type I IFN-inducing interactor of hnRNPM

To identify binding partners of hnRNPM involved in type I IFN induction, 28 putative interactors that were highly enriched or previously linked to antiviral immunity were transiently depleted from THP-1 cells by RNAi (targets and pathway annotations are labelled in Figure 2B). Next, the generated KD cell lines were stimulated with 5’ppp-dsRNA and pDNA to assess activation of RIG-I and cGAS signaling, respectively. We classified genes as relevant for cGAS or RIG-I signaling only if two shRNAs with high KD efficiencies impaired ISRE reporter activation. Although KD of the targeted interactors worked for both respective shRNAs, only KD of ELAVL1 markedly reduced the activation of the ISRE reporter after stimulation of cGAS or RIG-I in both shRNA expressing cell lines (Figures 3A, S3A–S3B).

**Figure 3:**
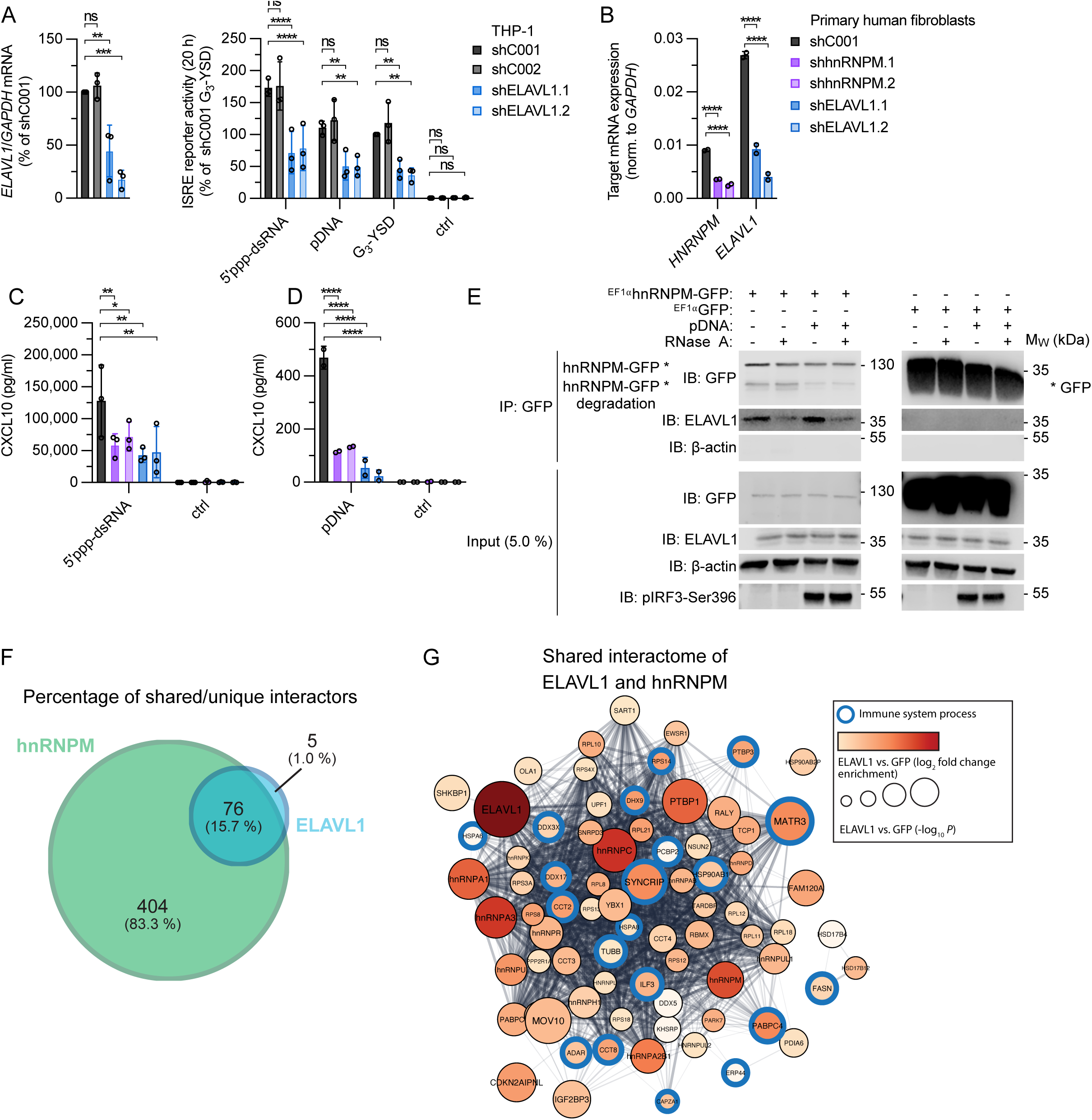
ELAVL1 interacts with hnRNPM to regulate type I IFN induction downstream of both cGAS and RIG-I. (A) Left panel: qPCR of *ELAVL1* in THP-1 cells expressing control shRNAs (shC001, shC002) or ELAVL1-specific shRNAs (shELAVL1.1, shELAVL1.2) (mean ± SD, one-way ANOVA, Dunnett’s multiple comparisons test), right panel: ISRE reporter activation 20 h after stimulation with 5’ppp-dsRNA (0.1 µg/ml), pDNA (0.1 µg/ml), or G_3_-YSD (0.5 µg/ml). ctrl, non-stimulated. (B) qPCR of *HNRNPM* and *ELAVL1* in human fibroblasts from adenoid tissue stably expressing control shRNA (shC001) or shRNAs against *HNRNPM* or *ELAVL1* (shRNA.1, shRNA.2). (C) CXCL10 ELISA with supernatants of the cells depicted in (B) 20 h after stimulation with 5’ppp-dsRNA (0.1 µg/ml). ctrl, non-stimulated. (D) CXCL10 ELISA with supernatants of the cells depicted in (B) 20 h after stimulation with pDNA (1 µg/ml). ctrl, non-stimulated. (E) GFP-specific beads were used to immunoprecipitate hnRNPM-GFP and GFP from lysates of non-stimulated or pDNA-stimulated (0.1 µg/ml, 3 h) THP-1 cells. If indicated, the IPs were treated with RNase A (100 µg/ml, 1.5 h). The IPs and 5.0% of the cleared cellular lysate used for IP (input) were analyzed by immunoblotting with the indicated antibodies. One representative experiment of two independent experiments is shown. (F) hnRNPM-GFP, ELAVL1-GFP, and GFP were immunoprecipitated from lysates of non-stimulated THP-1 cells and bound proteins were identified by LC-MS/MS. Detected proteins in hnRNPM and ELAVL1 precipitates were statistically compared to GFP as control using a two-sided Welch’s t-test (S0 = 0.1, permutation-based FDR < 0.05). The Venn diagram shows the absolute and relative proportions of unique and shared interactors of hnRNPM and ELAVL1. (G) Shared interaction network of hnRNPM and ELAVL1. Blue circles indicate proteins annotated with the gene ontology (GO) term immune system process (GO.0002376, FDR = 0.0034). For (A)–(D): mean ± SD, two-way ANOVA, Dunnett’s multiple comparisons test.

To evaluate the function of hnRNPM and ELAVL1 in primary cells, we performed an shRNA-mediated KD of hnRNPM or ELAVL1 in fibroblasts isolated from human adenoid tissue (Figures 3B–3D). In line with what was observed in THP-1 cells, fibroblasts depleted of hnRNPM or ELAVL1 secreted significantly less CXCL10, a surrogate marker for type I IFNs, after stimulation with 5’ppp-dsRNA or pDNA, indicating that the role of hnRNPM and ELAVL1 in RIG-I and cGAS signaling is not solely restricted to THP-1 but also relevant in primary and non-myeloid human cells.

It has been reported that hnRNPM shuttles from nucleus to cytoplasm after infection with SFV or CHIKV ^28^. Analysis of the subcellular localization showed that, under basal conditions, hnRNPM and ELAVL1 are predominantly localized in the nucleus (Figure S3C–S3D). Stimulation of cGAS or RIG-I did not alter the distribution of hnRNPM or ELAVL1 between nucleus and cytoplasm.

Next, we confirmed the interactions of hnRNPM and ELAVL1 by co-immunoprecipitation (co-IP) (Figure 3E). ELAVL1 interacted with hnRNPM in both the non-stimulated condition and after activation of cGAS. Co-precipitation of ELAVL1 with hnRNPM slightly decreased upon RNase A treatment, suggesting partially RNA-dependent interactions. We then asked whether hnRNPM and ELAVL1 are components of the same multiprotein complex. Therefore, we used AP-MS to also map the interactome of ELAVL1. To identify shared interactors of hnRNPM and ELAVL1, we compared the differential interactomes of hnRNPM and ELAVL1 (Figures 3F–3G; Table S1). Among the hnRNPM interactors, 15.7% were also detected in the interactome of ELAVL1, whereas 99.0% of the binding partners of ELAVL1 also interacted with hnRNPM, indicating that hnRNPM and ELAVL1 are part of the same multiprotein complex. We mapped the interactors shared by hnRNPM and ELAVL1 on the human protein-protein interaction network to identify a subnetwork of proteins that is shared by both baits (Figure 3G). Interestingly, several common interactors have previously been connected to immune system-related processes. Given this high enrichment of proteins related to immune system regulation, we speculate that other as yet undiscovered regulators of cGAS and RIG-I signaling exist within this network, whose functional evaluation is beyond the scope of this manuscript. In summary, we identified hnRNPM and its binding partner ELAVL1 as novel components of both the cGAS and RIG-I signaling pathways that non-redundantly promote the induction of type I IFNs.

### ELAVL1 non-redundantly promotes type I IFN induction and NF-κB activation elicited by both cGAS- and RIG-I

To further evaluate the role of ELAVL1, we targeted the *ELAVL1* gene locus in THP-1 cells by CRISPR-Cas9 with two different gRNAs (clones #1–2, ELAVL1 gRNA AN; clone 3, ELAVL1 gRNA AI) (Figure 4A). As expected, KO of ELAVL1 severely compromised ISRE reporter activation after stimulation of cGAS or RIG-I (Figure 4B). In contrast, activation of the ISRE reporter by recombinant IFNα was unaltered between wildtype (WT) and ELAVL1 KO cells, indicating that ELAVL1 does not influence IFNAR signaling (Figure 4B).

**Figure 4:**
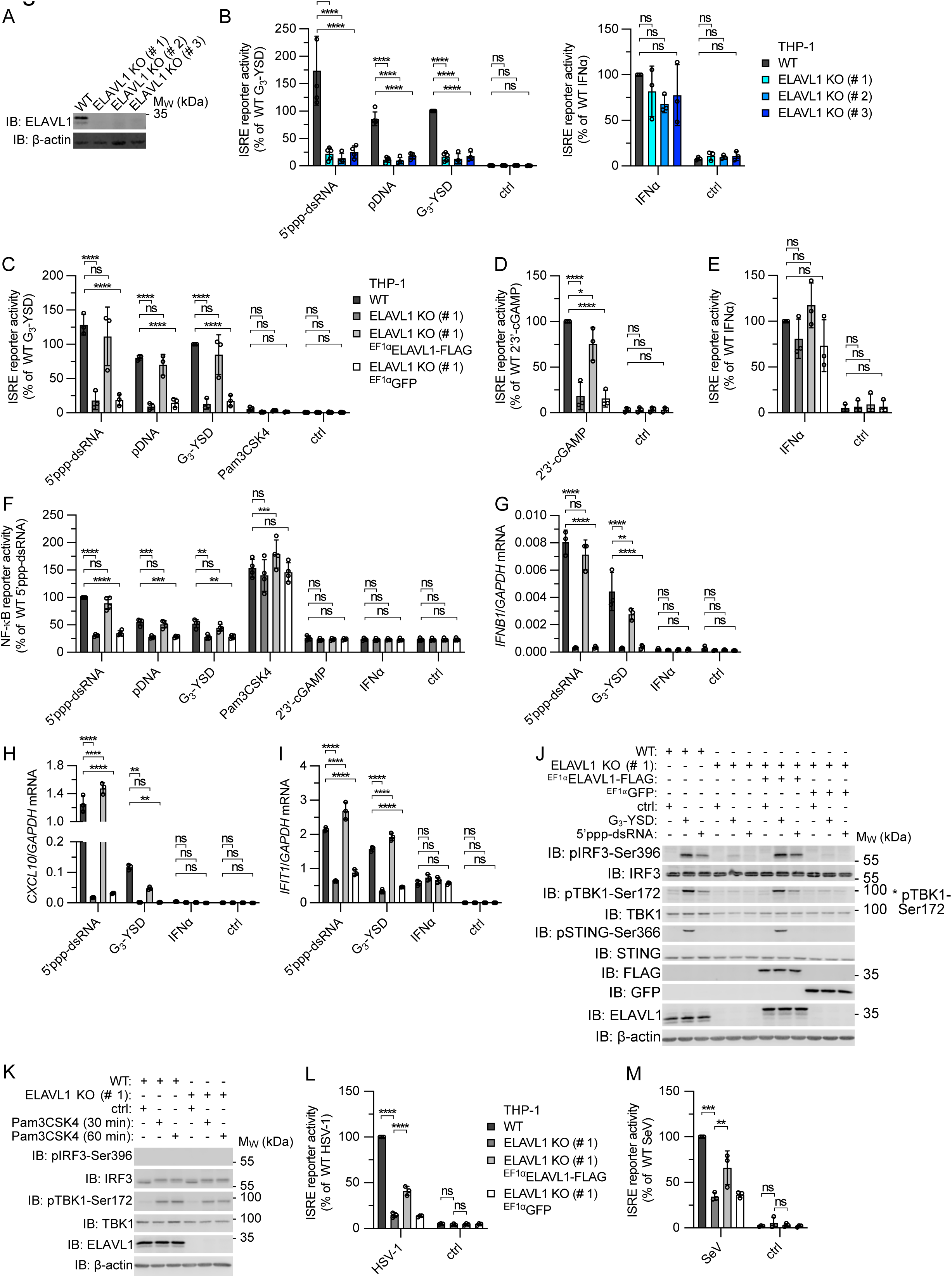
ELAVL1 promotes the cGAS- or RIG-I-mediated induction of type I IFNs and activation of NF-κB. (A) Immunoblot analysis of THP-1 WT and ELAVL1 KO cells using an ELAVL1-specific antibody. (B) ISRE reporter activation in the cells depicted in (A) 20 h after stimulation with 5’ppp-dsRNA (0.1 µg/ml), pDNA (0.1 µg/ml), G_3_-YSD (0.5 µg/ml), or IFNα (1000 U/ml). ctrl, non-stimulated. (C) ISRE reporter activation in THP-1 WT, ELAVL1 KO (#1), and ELAVL1 KO cells expressing ELAVL1-FLAG or GFP 20 h after stimulation with 5’ppp-dsRNA (0.1 µg/ml), pDNA (0.1 µg/ml), G_3_-YSD (0.5 µg/ml), or Pam3CSK4 (0.5 µg/ml). ctrl, non-stimulated. (D) ISRE reporter activation in the cells depicted in (C) 20 h after stimulation with 2’3’-cGAMP (10 µg/ml). ctrl, non-stimulated. (E) ISRE reporter activation in the cells depicted in (C) 20 h after stimulation with IFNα (1000 U/ml). ctrl, non-stimulated. (F) NF-κB reporter activation in the cells depicted in (C) 20 h after stimulation with 5’ppp-dsRNA (0.1 µg/ml), pDNA (0.1 µg/ml), G_3_-YSD (0.5 µg/ml), Pam3CSK4 (0.5 µg/ml), 2’3’-cGAMP (10 µg/ml), or IFNα (1000 U/ml). ctrl, non-stimulated. (G) qPCR of *IFNB1* in the cells depicted in (C) 6 h after stimulation with 5’ppp-dsRNA (0.1 µg/ml), G_3_-YSD (0.5 µg/ml), or IFNα (1000 U/ml). ctrl, non-stimulated. (H) qPCR of *CXCL10* in the cells depicted in (C) 6 h after stimulation with 5’ppp-dsRNA (0.1 µg/ml), G_3_-YSD (0.5 µg/ml), or IFNα (1000 U/ml). ctrl, non-stimulated. (I) qPCR of *IFIT1* in the cells depicted in (C) 6 h after stimulation with 5’ppp-dsRNA (0.1 µg/ml), G_3_-YSD (0.5 µg/ml), or IFNα (1000 U/ml). ctrl, non-stimulated. (J) Immunoblot analysis of the cells depicted in (C) 3 h after stimulation with G_3_-YSD (0.5 µg/ml) or 5’ppp-dsRNA (0.1 µg/ml). ctrl, non-stimulated. (K) Immunoblot analysis of THP-1 WT and ELAVL1 KO (#1) cells after stimulation with Pam3CSK4 (0.5 µg/ml). ctrl, non-stimulated. (L) ISRE reporter activation in the cells depicted in (C) 24 h after infection with HSV-1 (MOI 5). ctrl, non-stimulated. (M) ISRE reporter activation in the cells depicted in (C) 24 h after infection with SeV (MOI 1). ctrl, non-stimulated. For (B)–(I), (L)–(M) mean ± SD, two-way ANOVA, Dunnett’s multiple comparisons test. For (J)–(K): One representative experiment of at least two independent experiments is shown.

To exclude off-target effects, ELAVL1-FLAG was lentivirally expressed in THP-1 ELAVL1 KO clone #1. As expected, expression of ELAVL1-FLAG but not GFP rescued ISRE reporter activation in cells with ELAVL1-KO genetic background after stimulation of RIG-I, cGAS, or STING, demonstrating that ELAVL1 is a specific positive regulator downstream of these receptors (Figures 4C–4E). Reconstitution of ELAVL1 expression also re-established the RIG-I- and cGAS-dependent activation of the NF-κB reporter, while NF-κB signaling induced by TLR1/2 (Pam3CSK4) was ELAVL1-independent (Figure 4F). To evaluate the contribution of ELAVL1 to the activation of the respective pathways, we compared THP-1 ELAVL1 KO cells with RIG-I KO, MAVS KO, and cGAS KO cells. As expected, KO of RIG-I or MAVS abolished the 5’ppp-dsRNA-induced ISRE and NF-κB reporter activation, whereas KO of cGAS only disrupted the pDNA- or G_3_-YSD-triggered activation of these reporters (Figure S4A–S4B). As seen above, KO of ELAVL1 inhibited the cGAS- or RIG-I-induced ISRE reporter activity by approximate 5–8-fold, suggesting that ELAVL1 is not essential for type I IFN production but represents a potent enhancer. Although ELAVL1 is involved in different RNA-processing steps, ELAVL1-deficient cells did not display decreased viability (Figure S4C). In fact, THP-1 ELAVL1 KO monocytes were more viable after stimulation of RIG-I or cGAS. In addition, we found that KO of ELAVL1 blunted the expression of *IFNB1* and *CXCL10* mRNA following activation of cGAS or RIG-I (Figures 4G–4H). In line with this observation, secreted CXCL10 was also undetectable in the supernatants of ELAVL1 KO cells after 5’ppp-dsRNA or pDNA challenge (Figure S4D). Similar to what was observed for the ISRE reporter gene assay, the cGAS- or RIG-I-mediated induction of *IFIT1* mRNA was reduced but not completely abolished in THP-1 ELAVL1 KO cells (Figure 4I). By contrast, *IFIT1* mRNA expression induced by recombinant IFNα remained unchanged in ELAVL1 KO cells compared to WT, further demonstrating that IFNAR signaling is not influenced by ELAVL1 and highlighting the specific role of ELAVL1 in the cGAS and RIG-I signaling cascades (Figure 4I).

In addition, we found that the phosphorylation of IRF3, STING, and TBK1 upon activation of RIG-I and/or cGAS was ELAVL1-dependent (Figure 4J). Because the Pam3CSK4-induced NF-κB reporter activity was independent of ELAVL1, we also analyzed the TLR1/2-mediated pTBK1-Ser172 induction in THP-1 ELAVL1 KO cells. Interestingly, the Pam3CSK4-induced phosphorylation of TBK1 at Ser172 was not impaired by KO of ELAVL1 (Figure 4K). These data suggest that ELAVL1 specifically promotes TBK1 phosphorylation induced by MAVS- or STING-but not MyD88-dependent pathways and that ELAVL1 is able to discriminate TBK1 molecules in a PRR-dependent manner. We then analyzed the role of ELAVL1 during infection with DNA and RNA viruses. Similar to hnRNPM, ELAVL1 was required for both the HSV-1 and SeV-induced ISRE reporter activation (Figures 4L–4M).

Considering that the relative mRNA levels of *IFNB1*, *CXCL10*, and *IFIT1* were downregulated to different extents in ELAVL1 KO as compared to control cells, we hypothesize that ELAVL1 has a dual function in innate immunity. On the one hand, ELAVL1 modulates TBK1 and IRF3 activation, and, on the other hand, it stabilizes certain mRNA species as reported previously for *IFNB1* or certain ISGs ^38, 43^. Stabilization of *IFNB1* mRNA by ELAVL1 could have direct consequences for the detection of cytoplasmic nucleic acids, as type I IFNs themselves enhance the nucleic acid-sensing capacities of the cell in a positive feedback loop by upregulating the expression of first category nucleic acid receptors such as RIG-I or, to a lesser extent, cGAS. To quantify the role of ELAVL1 beyond *IFNB1*-dependent effects, we generated THP-1 cells lacking both IFNAR2 and ELAVL1. We hypothesized that inhibition of cGAS and RIG-I signaling in THP-1 IFNAR2 ELAVL1 double-KO cells indicates an active role of ELAVL1 in signal transduction. Although, as expected, we observed a reduction in ISRE activation for IFNAR2 KO cells when compared to WT, additional deficiency of ELAVL1 substantially inhibited both cGAS and RIG-I signaling compared to IFNAR2 single KO cells (Figure S5A). Similar observations were made after KD of hnRNPM in THP-1 IFNAR2 KO cells (Figure S5B).

Considering that ELAVL1 is an RNA-binding protein involved in different RNA-processing steps, we performed 3’-mRNA sequencing to determine whether KO of ELAVL1 impairs the overall expression of ISGs induced by IFNα. Here, we were unable to detect significant changes in the upregulation of ISGs, further indicating that ELAVL1 is decoupled from the IFNAR pathway and processing of ISG RNAs (Figure S5C; Table S2). Similarly, transcript levels of the main components of the cGAS and RIG-I signaling pathways were unchanged between ELAVL1 KO and THP-1 WT cells (Figure S5C; Table S2).

In summary, these data show that ELAVL1 promotes type I IFN expression and NF-κB signaling downstream of both RIG-I and cGAS-STING in a non-redundant fashion. We hypothesize that ELAVL1 and hnRNPM couple both signaling pathways by regulating a converging signaling step shared by both pathways.

### hnRNPM forms a multiprotein complex with ELAVL1, TBK1, and NF-κB p65

Our data suggest that hnRNPM and ELAVL1 are part of a protein complex that regulates the induction of type I IFNs downstream of both cGAS and RIG-I but upstream of IRF3 phosphorylation. Since we identified IKKβ as an interactor of hnRNPM (Figure 2B), we tested for interactions with structurally/functionally related antiviral signaling molecules such as IKKα, IKKε, or TBK1 (Figure 5A). In line with our AP-MS data, we observed specific co-precipitation of IKKβ with hnRNPM. Intriguingly, the IKK-related kinases TBK1 and IKKε as well as NF-κB p65 also selectively co-precipitated with hnRNPM. Interactions of hnRNPM with IKKβ, TBK1, IKKε, and NF-κB p65 were detectable in untreated and cGAS-activated cells. These interactions were independent of RNA since RNase A treatment did not affect precipitation efficacy. pTBK1-Ser172 and NF-κB p65 phosphorylated at Ser536 (NF-κB pp65-Ser536) were also enriched after IP of hnRNPM. Compared to untreated cells, cGAS activation increased hnRNPM-associated phosphorylation of TBK1 and NF-κB p65. Although IKKα shares 51% sequence identity with IKKβ ^44^, we were unable to detect binding of IKKα to hnRNPM, demonstrating that hnRNPM has a higher specificity for IKKε and IKKβ. Similarly, no interactions of hnRNPM with pIRF3-Ser396, IRF3, cGAS, and β-actin were detected. Although STING phosphorylated at Ser366 (pSTING-Ser366) was not detected in hnRNPM precipitates, low quantities of total STING co-immunoprecipitated with hnRNPM. To provide a second layer of specificity, we immunoprecipitated transiently expressed ELAVL1-FLAG from lysates of HEK293FT cells and screened for interactions of ELAVL1 with the hnRNPM-associated proteins (Figure 5B). As expected, endogenous hnRNPM co-immunoprecipitated with ELAVL1, with interactions slightly decreasing upon RNase A treatment. In line with our hypothesis that hnRNPM and ELAVL1 contribute to the same protein complex, also TBK1 and NF-κB p65 interacted with ELAVL1-FLAG in an RNase A-independent manner. By contrast, no interactions of ELAVL1 with IKKε and IKKβ were detectable. Although, we cannot exclude that hnRNPM forms a second ELAVL1-independent complex that harbors IKKε and IKKβ, we would rather interpret these data as the results of less robust interaction of ELAVL1 within the complex at our precipitation conditions.

**Figure 5:**
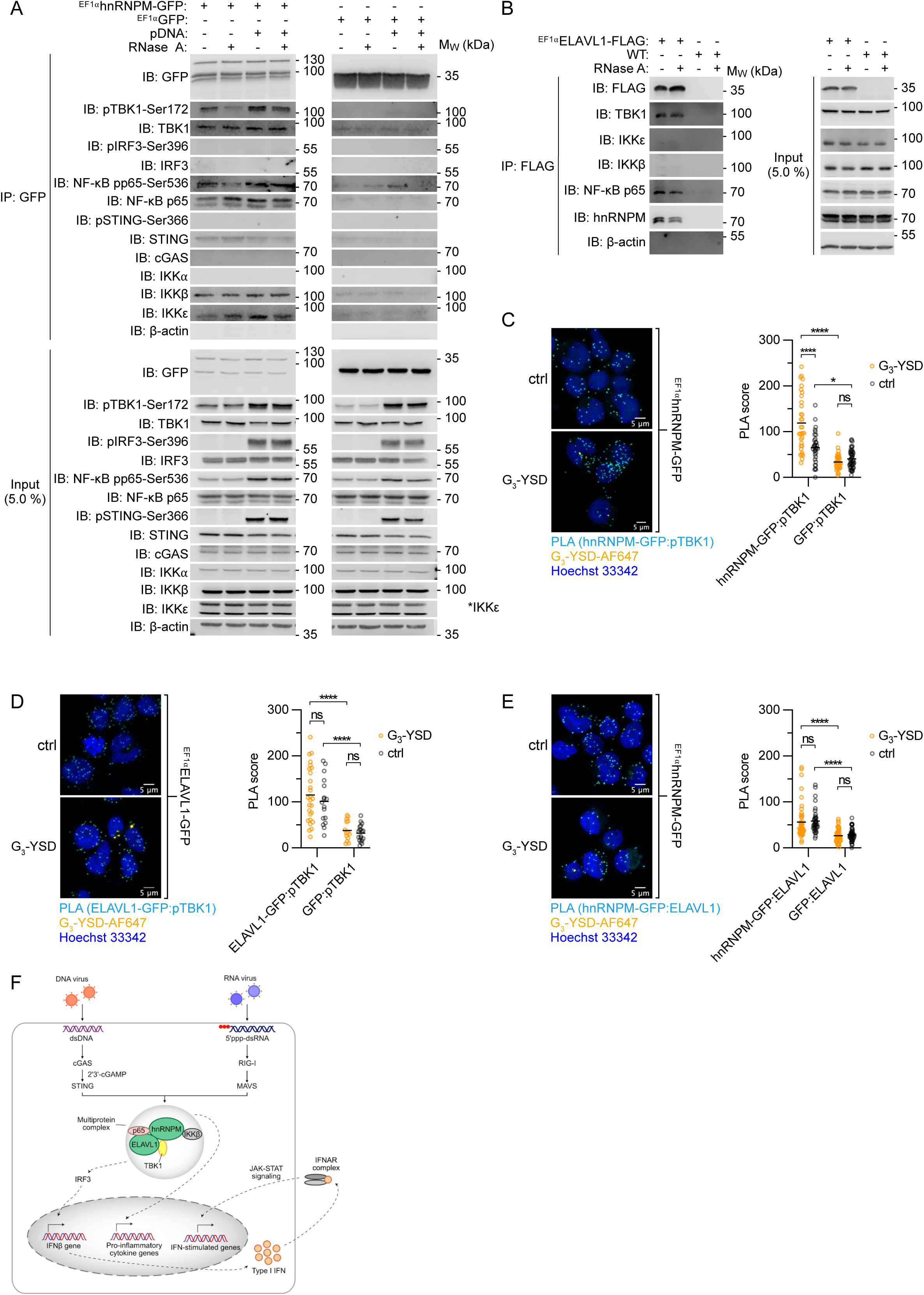
hnRNPM and ELAVL1 interact with TBK1 and NF-κB p65. (A) GFP-specific beads were used to immunoprecipitate hnRNPM-GFP and GFP (EF1α promotor) from lysates of non-stimulated or pDNA-stimulated (0.1 µg/ml, 3 h) THP-1 cells. If indicated, the IPs were treated with RNase A (100 µg/ml, 1.5 h). The IPs and 5.0% of the cleared cellular lysate used for IP (input) were analyzed by immunoblotting with the indicated antibodies. The IPs of hnRNPM-GFP and GFP were analyzed on the same membrane, with empty lanes removed. (B) FLAG tag-specific beads were incubated with lysates of WT or ELAVL1-FLAG-expressing HEK293FT cells. If indicated, the IPs were treated with RNase A (100 µg/ml, 1.5 h). The IPs and 5.0% of the cleared cellular lysate used for IP (input) were analyzed by immunoblotting with the indicated antibodies. (C) Differentiated THP-1 cells stably expressing hnRNPM-GFP or GFP were transfected with AF647-labelled G_3_-YSD (0.5 µg/ml, 4 h) or left non-stimulated (ctrl). Anti-GFP and anti-pTBK1-Ser172 monoclonal antibodies were used to analyze the interactions between hnRNPM-GFP/GFP and pTBK1-Ser172 *in cellulo* by PLA. Z stack images were recorded on a confocal microscope. Image sequences of hnRNPM-GFP-expressing cells are shown as maximum intensity projections (left panel) (GFP-expressing cells not shown). The PLA signals per cell in different focal planes were counted and the counted number of puncta per cell was defined as the PLA score (right panel). hnRNPM-GFP:pTBK1, interactions between hnRNPM-GFP and pTBK1-Ser172; GFP:pTBK1, interactions between GFP and pTBK1-Ser172. Blue, Hoechst 33342; yellow, G_3_-YSD-AF647; cyan, PLA. (D) The PLA was performed as described in (C) with differentiated THP-1 cells stably expressing ELAVL1-GFP or GFP (GFP-expressing cells not shown). (E) The PLA was performed as described in (C) with anti-GFP and anti-ELAVL1 monoclonal antibodies used to analyze the interactions between hnRNPM-GFP/GFP and ELAVL1 (GFP-expressing cells not shown). (F) Proposed model of how hnRNPM and ELAVL1 promote type I IFN induction upon stimulation of cGAS or RIG-I. For (A)–(B): One representative experiment of two independent experiments is shown. For (C)–(E): mean, two-way ANOVA, Tukey’s multiple comparisons test.

To determine whether the interactions between hnRNPM, ELAVL1, and pTBK1-Ser172 are direct or indirect, and to clarify whether these interactions also occur *in cellulo*, we performed proximity ligation assay (PLA) in differentiated THP-1 cells expressing hnRNPM-GFP, ELAVL1-GFP, or GFP, where positive signals occur only when the GFP-specific antibody is in close vicinity (≤ 40 nm) to the pTBK1-Ser172-specific antibody. It should be noted that a low PLA score was also detectable in GFP-expressing cells (Figures 5C–5D). However, significantly higher PLA scores were detected in cells expressing hnRNPM-GFP or ELAVL1-GFP (hnRNPM-GFP:pTBK1 or ELAVL1-GFP:pTBK1) as compared to GFP (GFP:pTBK1), indicating close interactions of hnRNPM and ELAVL1 with pTBK1-Ser172 in cells. For hnRNPM-GFP, the PLA score further increased upon cytosolic challenge with G_3_-YSD. Notably, interactions between ELAVL1 and pTBK1-Ser172 were also detectable under non-stimulated conditions. Assembly of the individual z stack images into 3D projections revealed that the majority of interactions was detectable in the cytoplasm (data not shown). We also confirmed close interaction between hnRNPM and ELAVL1 (Figure 5E). Here, we performed the PLA in hnRNPM-GFP-expressing cells using a combination of GFP- and ELAVL1-specific antibodies and observed a significantly higher PLA score for hnRNPM-GFP:ELAVL1 interaction as compared to GFP:ELAVL1. The interactions between hnRNPM and ELAVL1 were present in both cytoplasm and nucleus. In summary, our data link for the first time hnRNPM and ELAVL1 to active antiviral signaling proteins and provide evidence for the existence of a hitherto undiscovered protein complex that fuels the induction type I IFNs upon stimulation of cGAS or RIG-I (Fig. 5F).

## DISCUSSION

cGAS and RIG-I sense different ligands and signal via distinct adaptor proteins, but both their signaling cascades converge at the level of TBK1/IKK/IRF3/NF-κB p65, ultimately leading to the expression of type I IFNs. In the present study, we report the identification of a novel protein complex that promotes the expression of type I IFNs downstream of both cGAS and RIG-I. hnRNPM and ELAVL1 constitute two key components of the complex that non-redundantly control the phosphorylation of IRF3 after cytosolic challenge with ligands for cGAS or RIG-I. We could further show that the protein complex assembled by hnRNPM interacts with crucial signaling components downstream of cGAS and RIG-I, such as TBK1 and NF-κB p65, thereby merging both signaling pathways. It is conceivable that hnRNPM and ELAVL1 provide a scaffolding platform that stabilizes interactions and amplifies the phosphorylation of IRF3, thereby promoting type I IFN expression. Considering that hnRNPM and ELAVL1 share several binding partners, our AP-MS analysis strongly suggests that both proteins are part of the same multiprotein complex. This notion is further supported by our PLA data, demonstrating interactions between hnRNPM and ELAVL1 in cells. Interestingly, we also identified a number of proteins already connected to type I IFN induction as hnRNPM interactors, such as DDX3X, DHX9, DHX15, PRKDC, and ZC3HAV1. However, KD of these proteins did not significantly inhibit ISRE reporter activation after stimulation of cGAS or RIG-I, possibly due to cell type-/stimulus-specific effects or redundant functions. Similarly, other hnRNPs were dispensable for ISRE activation, indicating that hnRNPM may have a unique antiviral activity within the hnRNP family. Since a limited number of interactors were included in the RNAi screen, we suspect that other components of the multiprotein complex regulating cGAS and RIG-I signaling remain to be discovered, which was beyond the scope of this manuscript. It remains elusive why the interactions of hnRNPM with TBK1 and NF-κB p65 could not be captured by our AP-MS analysis. Low peptide ionization efficiencies of the prey proteins during MS analysis could provide a possible explanation.

Our biochemical and genetic analyses demonstrate that ELAVL1 specifically promotes the expression of type I IFNs and activation of NF-κB downstream of cGAS-STING and RIG-I and that ELAVL1 is functionally decoupled from IFNAR signaling. We demonstrate that ELAVL1 controls the phosphorylation of TBK1, IRF3, and STING after activation of cGAS and/or RIG-I. Functional decoupling of ELAVL1 from TLR1/2-dependent NF-κB signaling was also reflected by the unaltered Pam3CSK4-induced phosphorylation of TBK1 in THP-1 ELAVL1 KO cells. Considering that ELAVL1 drives the phosphorylation of TBK1 in a PRR-dependent manner, we infer that TBK1 molecules are recruited to distinct complexes which can be distinguished by ELAVL1. How ELAVL1 discriminates different TBK1 populations will be the subject of future research. In general, pre-formed protein complexes are advantageous for the cell to rapidly orchestrate a directed immune response to exogenous triggers while providing improved means for spatiotemporal control due to the local accessibility of signaling proteins. Interestingly, although hnRNPM and ELAVL1 are both described as RNA-binding proteins, co-precipitation of TBK1 and NF-κB p65 with hnRNPM/ELAVL1 was RNA-independent, suggesting that the interactions are mediated by protein domains and not by structural RNA sequences such as lncRNA. Based on the PLA, we conclude that the heterotypic interactions of hnRNPM and ELAVL1 with pTBK1-Ser172 are predominantly localized in the cytoplasm, suggesting that the minor cytoplasmic rather than the predominating nuclear fraction of the hnRNPM-ELAVL1 complex possesses the antiviral activity.

ELAVL1 is a well-studied RNA-binding protein composed of three RRMs and known to stabilize different mRNAs by binding to AU-rich elements in the 3’-UTR. Consequently, ELAVL1 has been implicated in several cellular processes ranging from development to angiogenesis and has also been linked to inflammatory diseases and cancer ^45, 46^. Considering the role of ELAVL1 in transcript stabilization, an indirect effect on cGAS and RIG-I signaling by mRNA stabilization is thinkable. However, we could not detect decreased expression of the main components of the cGAS and RIG-I signaling pathways in our 3’-mRNA sequencing analysis, and the ISRE reporter activity induced by cGAS or RIG-I was also severely dampened in THP-1 cells with ELAVL1 and IFNAR2 double-KO genetic background, demonstrating that ELAVL1 promotes signaling upstream of IFNAR. Additionally, ELAVL1 and hnRNPM interact with TBK1 and NF-κB p65. Therefore, besides a possible involvement in mRNA stabilization, our data provide evidence for a direct role of ELAVL1 and hnRNPM innate antiviral signaling.

Notably, we find that hnRNPM/ELAVL1 promote an innate immune response against both HSV-1 and SeV. A broad biological relevance of the hnRNPM complex is further highlighted by the observation that KD of hnRNPM or ELAVL1 also impairs both cGAS and RIG-I signaling in primary human fibroblasts. Based on our data, it is conceivable that hnRNPM and ELAVL1 represent attractive new targets for the treatment of autoinflammatory disorders resulting from the pathological activation of cGAS, STING, or RIG-I. New therapeutics would not only be a particularly promising strategy for the treatment of monogenetic type-I interferonopathies but also harbor great potential for the growing number of degenerative diseases that have been associated with cGAS-STING activation, such as Huntington disease, Parkinson disease, and amyotrophic lateral sclerosis ^47, 48^. Ultimately, a more detailed understanding of this newly defined protein complex will need to be elucidated in futures studies. Nonetheless, the identification of hnRNPM and ELAVL1 as important components of cGAS-STING and RIG-I-MAVS signaling will provide more insight into the interaction of pathogens with our innate immune system, such as the previously observed interaction between SARS-CoV1 ORF3b and hnRNPM.

## MATERIAL AND METHODS

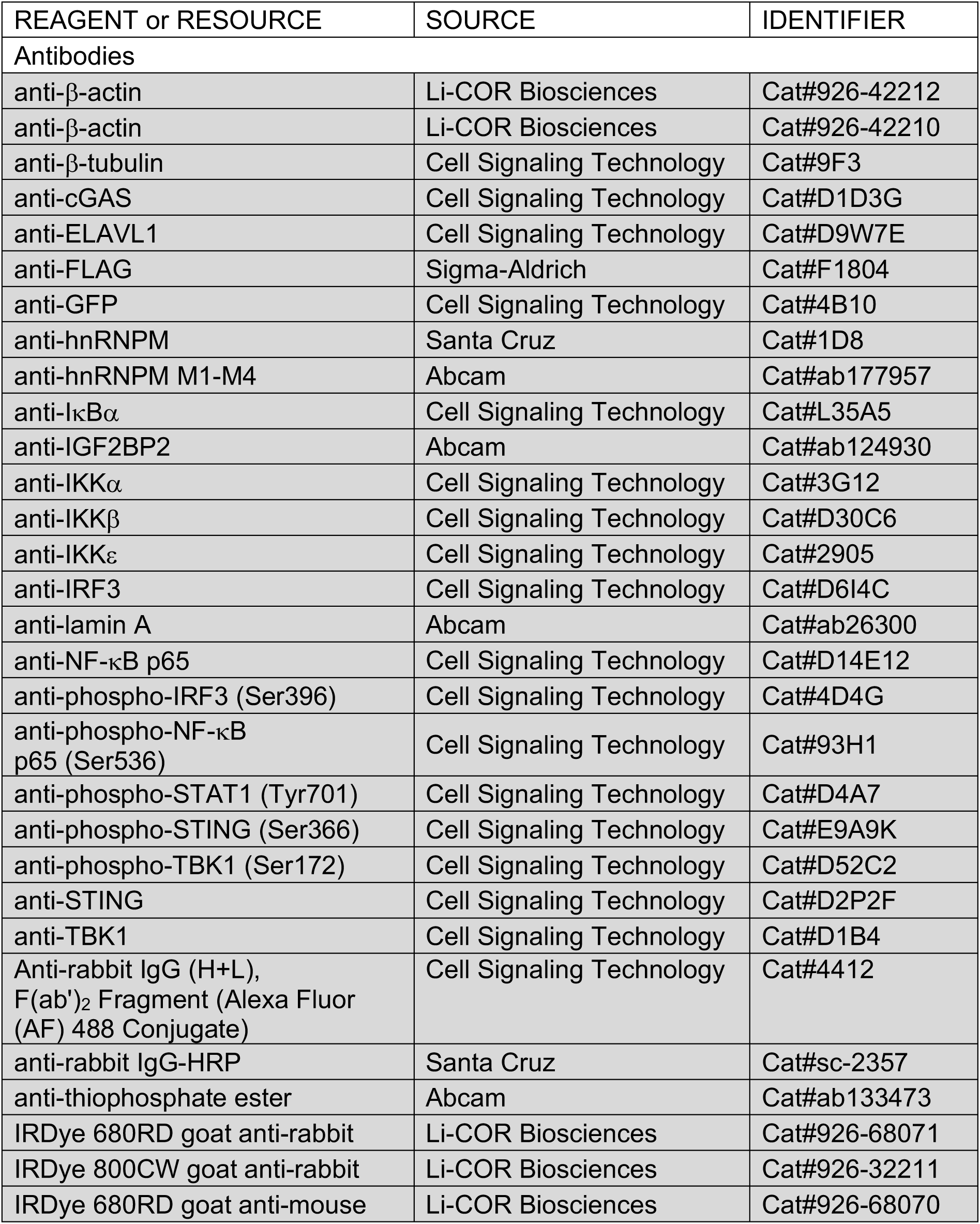
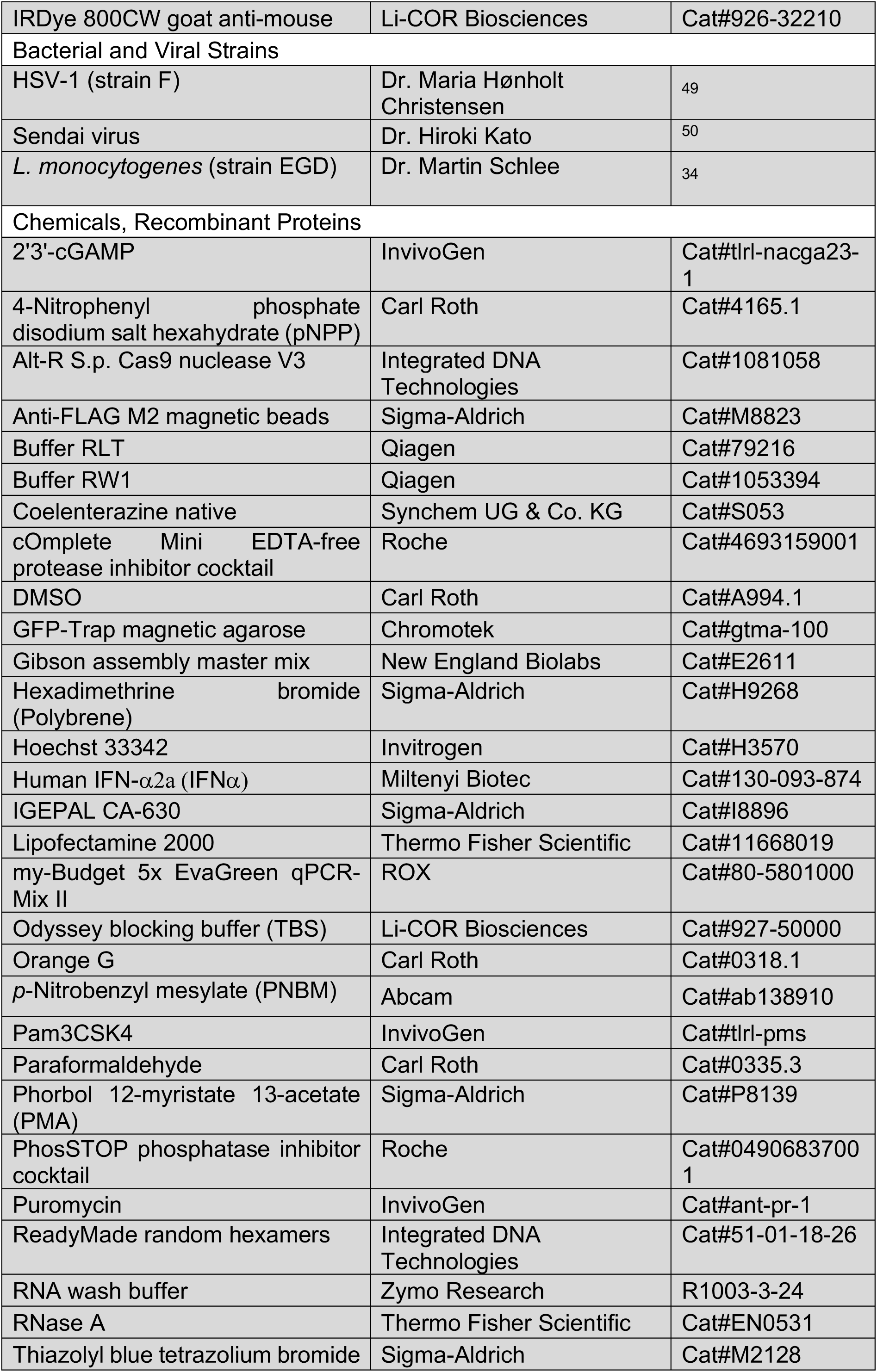
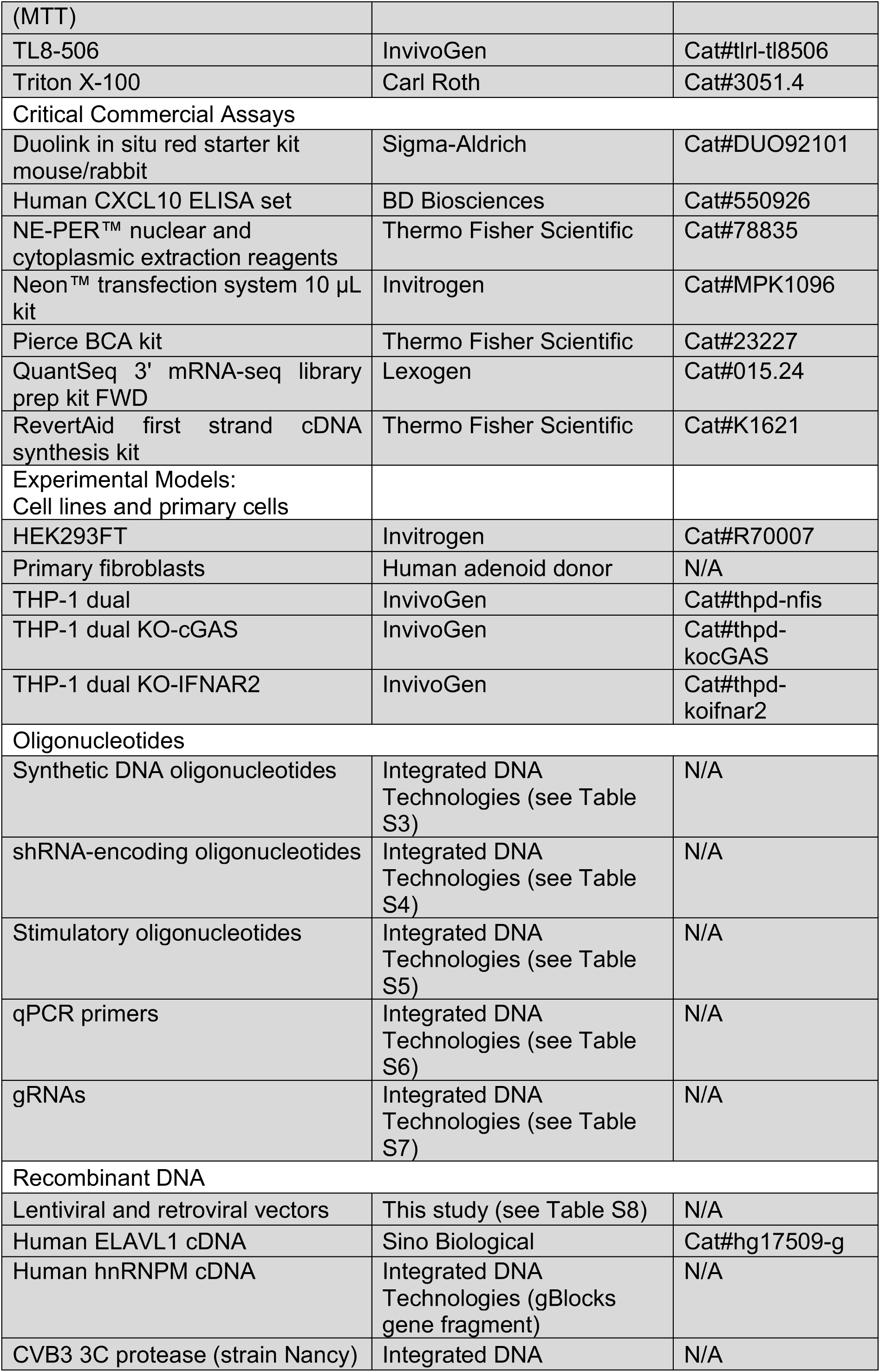
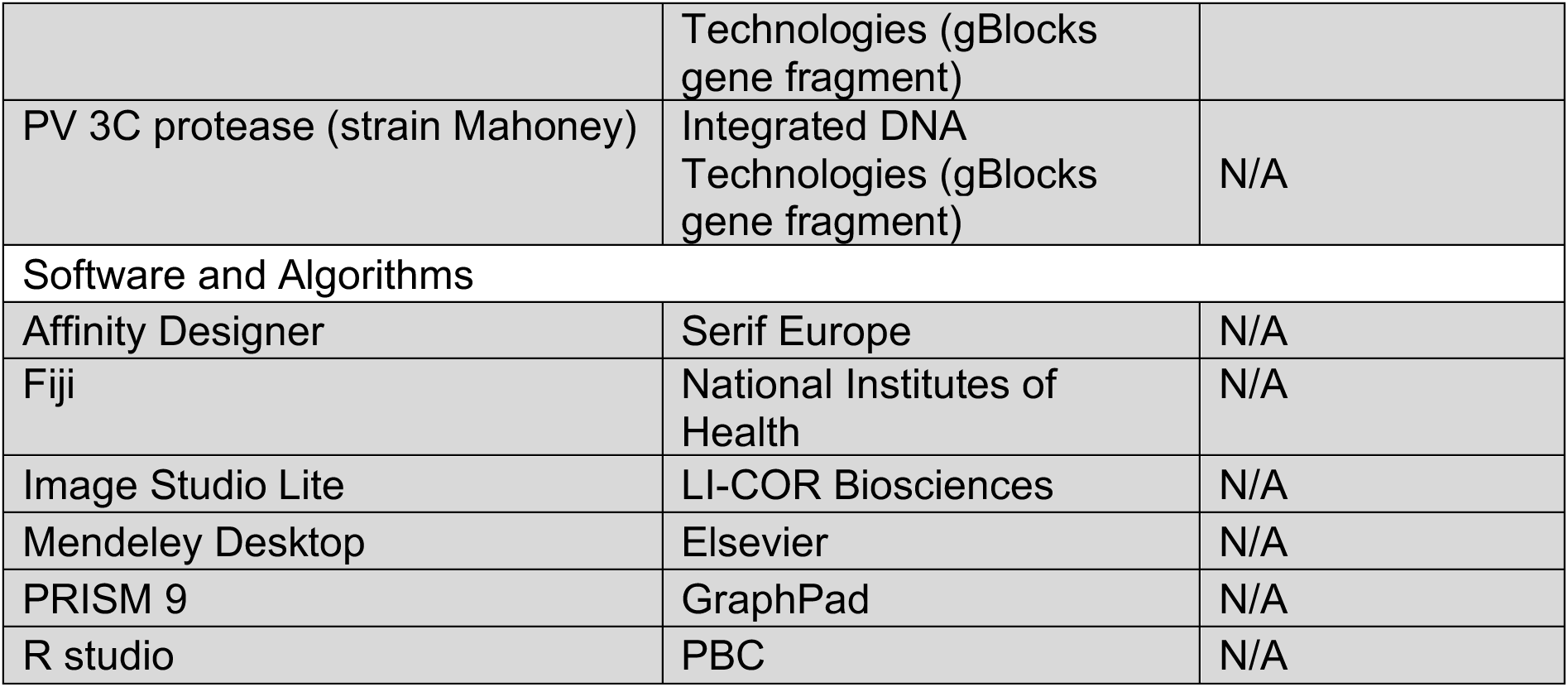
RESOURCES TABLE

## LEAD CONTACT AND MATERIALS AVAILABILITY

Further information and requests for resources and reagents should be directed to and will be fulfilled by the Lead Contact, Martin Schlee.

## EXPERIMENTAL MODEL AND SUBJECT DETAILS

### Ethics statement

The studies involving primary human fibroblasts were approved by the local ethics committee (Ethics committee of the Medical Faculty of the University of Bonn). Written informed consent was provided by the donors.

### Cell culture

THP-1 cells were cultured in RPMI 1640 (10% FCS, 100 U/ml penicillin, 100 mg/ml streptomycin) at 37 °C. HEK293FT cells were cultured in DMEM (10% FCS, 100 U/ml penicillin, 100 mg/ml streptomycin) at 37 °C. Primary fibroblasts were isolated from human adenoid tissue and cultured in DMEM at 37 °C.

## METHOD DETAILS

### Transient Expression of 3C proteases of PV and CVB3

10^5^ HEK293FT cells were seeded in 400 µl DMEM in 24-well plates. On the following day, the medium was refreshed and the cells were transiently transfected with pLenti^EF1α^ encoding hnRNPM-GFP or GFP together with pBluescript (control vector)/pRP^CMV^-CVB3 3C-protease-FLAG/pRP^CMV^-PV 3C-protease-FLAG (200 ng each) using 1.0 µl Lipofectamine 2000 (1 µg/µl) according to the manufacturer’s instructions. After 48 h, GFP expression was analyzed using an Axio Vert.A1 fluorescence microscope (Zeiss).

### Generation of cleared cellular lysates

Routinely, 3.6 × 10^5^ THP-1 cells were washed with PBS and lysed in 25 µl RIPA buffer (150 mM NaCl, 50 mM Tris/HCl, 1.0% (v/v) Triton X-100, 0.5% (w/v) sodium deoxycholate, 0.1% (w/v) SDS, pH 8.0) supplemented with protease and phosphatase inhibitor cocktail on ice for 20 min. Cellular debris were removed by centrifugation and the supernatant was isolated as the cleared cellular lysate. Following, 25 µl 2× Laemmli buffer (240 mM Tris/HCl, 8% (w/v) SDS, 40% (v/v) glycerol, 40 mM DTT, Orange G, pH 6.8) were added. Then, the samples were incubated at 95 °C for 5 min and subjected to immunoblot analysis.

### Protein quantification

Protein concentrations were determined using the Pierce bicinchoninic acid assay (BCA) kit (Thermo Fisher Scientific) according to the supplier’s protocol.

### Cloning techniques

The plasmids listed in Table S8 were cloned using Gibson assembly or traditional cloning and the primers listed in Tables S3–S4.

### Cloning of pLKO.1 KD vectors

shRNA sequences were from the MISSION shRNA library (Sigma-Aldrich) or designed with the siRNA Wizard Software (InvivoGen). To clone shRNA-expressing vectors, the pLKO.1 shC001 vector (Sigma-Aldrich) was digested with *Age*I and *Eco*RI (Thermo Fisher Scientific) and ligated with the shRNA-encoding, annealed DNA oligonucleotides (Integrated DNA Technologies) (Table S4). The ligation product was transformed into *E. coli* Stbl3 and positive colonies were identified by Sanger sequencing.

### Lentiviral expression

Lentiviral vectors with pLenti-blasticidin-EF1α and pLVX-puromycin-EF1α backbones (1.6 µg) (Table S8) were co-transfected into HEK293FT cells with pRSV-Rev (0.4 µg), pMDLg/pRRE (1.1 µg), and pMD2.G (0.6 µg) (6-well format) using standard CaPO_4_ precipitation. The shRNA-expressing pLKO.1 vectors (1 µg) (Table S8) were co-transfected into HEK293FT cells with psPAX2 (0.75 µg) and pMD2.G (0.25 µg). After 72 h, the virus-containing supernatants were harvested and passed through 0.45 µm filters. The viral supernatants were spiked with 5 µg/ml polybrene (Sigma-Aldrich) and then the indicated THP-1 cells were spin-infected at 32 °C at 600 ×g for 60 min. Primary human fibroblasts were conventionally infected with the polybrene-containing virus solution. After 24 h, the medium was refreshed and transduced cells were selected with 2 µg/ml puromycin (InvivoGen) for 3 days or sorted by FACS to isolate cell populations with comparable GFP expression levels. Polyclonal cell populations were used for subsequent experiments.

### qPCR

For isolation of total RNA, cells were resuspended in 350 µl RLT buffer (Qiagen) and then 350 µl 70% (v/v) ethanol was added to the mixture. The samples were loaded onto Zymo Spin IIICG columns (Zymo Research) and washed sequentially with 350 µl buffer RW1 (Qiagen) and 350 µl Zymo RNA wash buffer (Zymo Research). The RNA was eluted in RNase-free H_2_O. If required, samples were treated with DNase I (Thermo Fisher Scientific) according to the manufacturer’s protocol to remove residual genomic DNA. cDNA was synthesized using random hexamers (Integrated DNA Technologies) and the RevertAid first strand cDNA synthesis Kit (Thermo Fisher Scientific) according to the supplier’s instructions. qPCR was performed with my-Budget 5x EvaGreen qPCR-Mix II (ROX) on a QuantStudio 5 Real-Time PCR system device (Thermo Fisher Scientific) (qPCR primers listed in Table S6).

### Stimulatory nucleic acids

Annealed template DNA oligonucleotides (Table S5) were used to generate 5’ppp-dsRNA by *in vitro* transcription as described before ^51^. G_3_-YSD was prepared by hybridizing the single-stranded DNA oligonucleotides G_3_-YSD fwd and G_3_-YSD rev (Table S5). The oligonucleotides were mixed 1:1 in 1× NEBuffer 2, incubated at 95 °C for 5 min, and annealed by decreasing the temperature by 1 °C/min to 8 °C. pBluescript (pDNA) was isolated from transformed *E. coli* Stbl3 using the PureLink HiPure plasmid midiprep kit (Thermo Fisher Scientific).

### Cell stimulation

Stimulation experiments were routinely performed in 96-well plates in duplicates. Per well, 6 × 10^4^ cells THP-1 cells or 5 × 10^3^ primary human fibroblasts were seeded in 150 µl medium. 5’ppp-dsRNA (0.1 µg/ml), pDNA (0.1 µg/ml; 1.0 µg/ml), or G_3_-YSD (0.5 µg/ml) were transfected using Lipofectamine 2000. Per well, 0.5 µl Lipofectamine 2000 (1 µg/µl) and the RNA or DNA stimuli were diluted in Opti-MEM in a total volume of 25 µl each. Following, both mixtures were combined, incubated for 5 min at room temperature, and added to the cells. 2’3’-cGAMP (10 µg/ml), TL8-506 (1.0 µg/ml), Pam3CSK4 (0.5 µg/ml), and IFNα (1000 U/ml; 5000 U/ml) were diluted to the desired concentrations in 50 µl Opti-MEM and directly added to the cells. In the non-stimulated condition (ctrl), 50 µl Opti-MEM was added. After the indicated times, the supernatants were harvested and used for downstream assays. If assays required larger formats, the experimental parameters were adjusted proportionally, considering the surface area of the well.

For viral infections, 3 x 10^5^ THP-1 cells were seeded in 300 µl RPMI and infected with HSV-1 or SeV at the indicated MOI. After 24 h, activation of the ISRE reporter was determined. For infection with *L. monocytogenes*, 3 x 10^5^ THP-1 cells were seeded in antibiotics-free medium and incubated with *Listeria* at MOI 1 for 2 h. Following, the cells were washed with medium devoid of antibiotics and resuspended in medium containing 50 µg/ml gentamicin. After 24 h, activation of the ISRE reporter was determined.

### Detection of cytokines

Secretion of CXCL10, IL-6, and TNF-⍺ was analyzed with a commercial ELISA assay kit (BD Biosciences) according to the manufacturer’s instructions.

### MTT assay

Metabolic activity was determined by MTT assay and used as a surrogate parameter for cellular viability. 6 × 10^4^ cells were cultured in 100 µl medium supplemented with the yellow tetrazole substrate compound MTT (1 mg/ml) for 1 h. The reaction was stopped by adding 100 µl SDS solution (10% (w/v)) and the absorbance at λ = 595 nm was measured after dissolution of formazan crystals.

### Quantification of ISRE and NF-κB reporter activation in THP-1 dual cells

THP-1 dual cells stably express a Gaussia luciferase gene under the control of an ISG54 minimal promotor fused to five ISREs, and a secreted alkaline phosphatase (SEAP) gene under the control of an IFNβ minimal promotor in conjunction with five NF-κB consensus transcriptional response elements and three c-Rel-binding sites. To determine ISRE reporter gene activity, 30 µl cell culture supernatant and 30 µl coelenterazine solution (1 µg/ml in H_2_O) were mixed and the luminescence signal was measured using an EnVision 2104 Multilabel Reader Device (PerkinElmer). For the NF-κB reporter, 40 µl cell culture supernatant were incubated with 40 µl pNPP substrate buffer (100 mM NaCl, 100 mM Tris/HCl, 5 mM MgCl_2_, 10 mg/ml pNPP, pH 9.5) for 30–60 min and reporter gene activation was determined by measuring the absorbance at α = 405 nm.

### Generation of nuclear and cytoplasmic extracts

1.86 × 10^6^ cells were lipofected with 0.5 µg/ml G_3_-YSD or 0.1 µg/ml 5’ppp-dsRNA, or left non-stimulated. After 3 h, cells were harvested, washed with PBS, and the NE-PER nuclear and cytoplasmic extraction kit (Thermo Fisher Scientific) was used to isolate nuclear and cytoplasmic fractions according to the supplier’s protocol. Protein concentrations were determined by BCA assay (Thermo Fisher Scientific) and 5–10 µg total protein were analyzed by immunoblotting.

### Immunoblotting

Samples were separated by discontinuous, denaturing SDS-PAGE using 3% SDS-PAGE stacking gels and 10% SDS-PAGE resolving gels. After separation, proteins were transferred to 0.45 µm nitrocellulose membranes. If multiple targets were analyzed, equal quantities of the protein samples were resolved on different SDS-PAGE gels and membranes were probed sequentially. Ponceaus S staining was used to examine the success of the protein transfer. Membranes were destained with TBS containing 0.1 % (v/v) Tween 20, blocked with 5% BSA-PBS containing 0.1 % (v/v) Tween 20 for 1 h, and incubated with the indicated primary (1:500–1:1,000) and secondary antibodies (1:12,500). Protein signals were recorded on an Odyssey Fc near-infrared imaging system device (LI-COR Biosciences) and the signal intensities were analyzed using the Image Studio Lite software (LI-COR Biosciences).

### AP-MS analysis

THP-1 cells expressing hnRNPM-GFP, ELAVL1-GFP, or GFP were lysed in TAP lysis buffer (100 mM NaCl, 50 mM Tris/HCl, 1.5 mM MgCl_2_, 5% (v/v) glycerol, 0.5% (v/v) IGEPAL CA-630, pH 7.5) supplemented with protease and phosphatase inhibitor cocktail on ice for 20 min, followed by a short sonication burst (15 s, amplitude 90%, cycle 1, VialTweeter Sonication Device) (quadruplicates of each cell line). 25 µl equilibrated GFP-Trap magnetic agarose beads (Chromotek) were added to the clarified lysates (1.5 mg total protein per replicate) and incubated under rotation at 4°C overnight. The beads were washed with TAP lysis buffer (3×, 700 µl) and TAP wash buffer (100 mM NaCl, 50 mM Tris/HCl, 1.5 mM MgCl_2_, 5% (v/v) glycerol, pH 7.5) (3×, 700 µl) and stored at -80 °C for further processing. Enriched proteins were denatured with 40 µl U/A buffer (8 M urea, 100 mM Tris/HCl, pH 8.5), reduced with 10 mM DTT (30 min, 25°C), alkylated with 5.5 mM IAA (20 min, 25°C, in the dark) and digested by subsequent addition of 0.5 µg LysC (3 h, 25 °C; WAKO Chemicals) and 0.5 µg Trypsin (18 h, 25 °C; Promega) in ABC buffer (50 mM NH_4_HCO_3_, pH 8.0). Peptide purification on StageTips with three layers of C18 Empore filter discs (3M) and subsequent mass spectrometry analysis was performed as described previously ^52, 53^. Briefly, purified peptides were loaded on a 50 cm reverse-phase analytical column (75 µm diameter, 60 °C; ReproSil-Pur C18-AQ 1.9 µm resin; Dr. Maisch) and separated using an EASY-nLC 1200 system (Thermo Fisher Scientific). For peptide separation, a 120 min gradient with a flow rate of 300 nl/min and a binary buffer system consisting of buffer A (0.1% formic acid in H_2_O) and buffer B (80% acetonitrile, 0.1% formic acid in H_2_O) was used: 5–30% buffer B (95 min), 30–95% buffer B (10 min), wash out at 95% buffer B (5 min), decreased to 5% buffer B (5 min), and kept at 5% buffer B (5 min). Eluting peptides were directly analyzed on a Q-Exactive HF mass spectrometer equipped with a nano-electrospray source (Thermo Fisher Scientific). Spray voltage was set to 2.4 kV, funnel RF level at 60, and heated capillary at 250 °C. Data-dependent acquisition included repeating cycles of one MS1 full scan (300–1650 m/z, R = 60 000 at 200 m/z) at an ion target of 3 × 10^6^ with an injection time of 20 ms, followed by 15 MS2 scans of the highest abundant isolated and higher-energy collisional dissociation (HCD) fragmented peptide precursors (R = 15 000 at 200 m/z). For MS2 scans, collection of isolated peptide precursors was limited by an ion target of 1 × 10^5^ and a maximum injection time of 25 ms. Isolation and fragmentation of the same peptide precursor was eliminated by dynamic exclusion for 20 s. The isolation window of the quadrupole was set to 1.4 m/z and HCD was set to a normalized collision energy of 27%. RAW files were processed with MaxQuant (version 1.6.17.0) using the standard settings and label-free quantification (iBAQ; LFQ, LFQ min ratio count 1, normalization type none) enabled. Spectra were searched against forward and reverse sequences of the reviewed human proteome including isoforms (UniprotKB, release 08.2020) and GFP by the built-in Andromeda search engine ^54^.

### Statistical analysis of AP-MS data

The output of MaxQuant was analyzed with Perseus (version 1.6.14.0), R (version 4.0.2), and RStudio (version 1.3.1073) ^55^. Detected protein groups identified as known contaminants, reverse sequence matches, or only identified by site were excluded from the analysis. Following log_2_ transformation, the iBAQ intensity of each protein group in a given sample was normalized to correct for technical variation by subtracting a sample-specific normalization factor (NF_j_) based on the median iBAQ intensity of all protein groups per sample (median_j_):

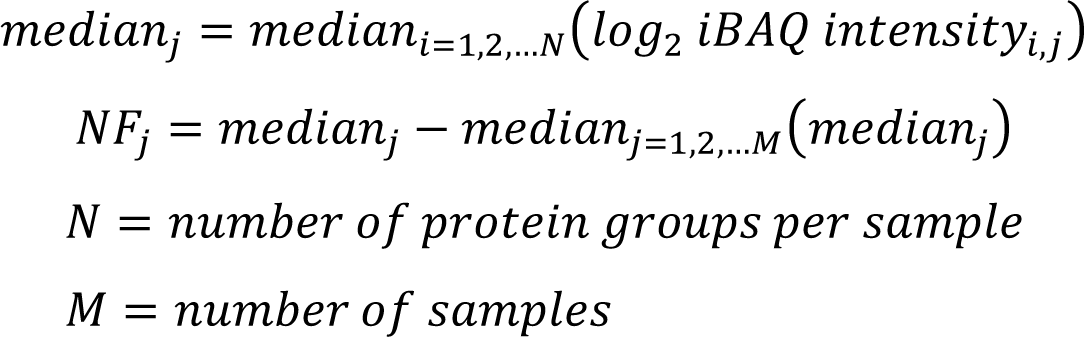

Following normalization, proteins without quantification in at least three replicates of one condition were removed and missing values were imputed for each replicate individually by sampling values from a normal distribution calculated from the original data distribution (width = 0.3 × s.d., downshift = -1.8 × s.d.). Differentially enriched protein groups between the conditions were identified via two-sided Welch’s t-tests (S0 = 0.1) corrected for multiple hypothesis testing applying a permutation-based FDR (FDR < 0.05, 250 randomizations). Protein groups that were not quantified in at least three replicates in the ELAVL1 or hnRNPM condition when compared to GFP or when comparing ELAVL1 to hnRNPM were removed for statistical testing. STRING-based analysis of hnRNPM or hnRNPM-ELAVL1 shared interactors was performed in Cytoscape (version 3.8.2) using the stringApp (version 1.6.0) in combination with a confidence cutoff of 0.2 for considering functional connections between interactors, an MCL inflation parameter of 3 for clustering and a Benjamini-Hochberg-adjusted FDR of < 0.05 to select significantly enriched pathway annotations.

### Co-IP

Prior to IP, THP-1 cells were transduced to express lentiviral particles encoding hnRNPM-GFP or GFP and populations with comparable GFP expression were isolated by FACS. 7.5 × 10^6^ cells were seeded in 7.5 ml RPMI and lipofected with 0.1 µg/ml pDNA to activate cGAS or treated with Opti-MEM. After incubation for 3 h, the cells were washed with PBS and lysed in 700 µl TAP lysis buffer supplemented with protease and phosphatase inhibitor cocktail for 20 min on ice. Using a short sonication burst, the nuclei were disrupted and the lysate was cleared from debris by centrifugation. 700 µg to 800 µg total protein were incubated with 12.5 µl equilibrated GFP-Trap magnetic agarose beads (Chromotek) on a rotating wheel at 4 °C overnight. If RNase A treatment was performed, the beads were washed with TAP wash buffer (3×, 700 µl), resuspended in 300 µl TAP wash buffer supplemented with protease inhibitor, phosphatase inhibitor and 100 µg/ml RNase A (Thermo Fisher Scientific), and incubated under rotation at 4 °C for 90 min. Following, the beads were washed sequentially with TAP lysis buffer (3×, 700 µl) and TAP wash buffer (3×, 700 µl) and bound proteins were eluted by incubating the beads for 10 min at 95 °C with 2× Laemmli buffer. Input loading controls (5% of total protein used for IP) and eluates were then subjected immunoblot analysis.

To immunoprecipitate ELAVL1, 2.5 × 10^6^ cells HEK293FT cells were seeded per 10-cm plate. On the next day, the medium was replaced and the cells were transfected with 20 µg pLVX^EF1α^-ELAVL1-FLAG using lipofectamine 2000 or left untreated. After 72 h, cleared cellular lysates were generated as described above. Per IP, 1 mg total protein was incubated with 50 µl equilibrated anti-FLAG M2 magnetic beads on a rotating wheel at 4 °C overnight. Unspecific protein binding to the beads incubated with lysate of WT cells was quenched with 4 ng/µl 3× FLAG peptide. Following, the beads were washed with TAP wash buffer (3×, 700 µl), resuspended in 300 µl TAP wash buffer supplemented with protease inhibitor, phosphatase inhibitor and, if indicated, 100 µg/ml RNase A. The samples were incubated under rotation at 4 °C for 90 min and subsequently the beads were washed sequentially with TAP lysis buffer (3×, 700 µl) and TAP wash buffer (3×, 700 µl). Bound proteins were eluted by incubating the beads for 30 min at 4 °C with 100 µl TAP wash buffer supplemented with 150 ng/µl 3× FLAG peptide. After magnetic separation, the eluates were incubated with Laemmli buffer at 95 °C for 5 min and subjected immunoblot analysis. 5% of the total protein used for IP were analyzed as input loading controls.

### CRISPR-Cas9-mediated KO cell line generation

The Alt-R CRISPR-Cas9 system (Integrated DNA Technologies) was used according to the supplier’s instructions to disrupt the genes encoding for ELAVL1, RIG-I, and MAVS in THP-1 cells. gRNAs (Table S7) were designed using the Alt-R HDR design tool (Integrated DNA Technologies). Briefly, crRNA:tracrRNA hybrids were formed by mixing 0.5 µl crRNA (200 µM), 0.5 µl tracrRNA (200 µM), and 1.28 µl IDTE buffer, followed by incubating the mixture for 5 min at 95 °C and gradual cooling to room temperature. Afterwards, 1.25 µl crRNA:tracrRNA hybrid was mixed with 1.25 µl diluted Cas9, which was prepared by mixing 0.5 µl resuspension buffer R (Neon transfection system 10 µl kit, Thermo Fisher Scientific) with 0.75 µl Cas9 enzyme, and incubated at room temperature for 20 min. 1.25 × 10^6^ THP-1 cells were washed with PBS, resuspended in 22.5 µl resuspension buffer R. 18 µl cell suspension was mixed with 4 µl electroporation enhancer (10.8 µM) (Integrated DNA Technologies) and 2 µl crRNA:tracrRNA:Cas9 RNP complex. Electroporations were performed on a Neon electroporation system device (Thermo Fisher Scientific) at 1600 V with 3 pulses and 10 ms pulse width using a 10 µl Neon tip. Cells were cloned by limiting dilution and target KOs were verified was immunoblotting and/or Sanger sequencing.

### 3’-mRNA sequencing

THP-1 WT and ELAVL1 KO (clone #1) cells were stimulated with 1000 U/ml IFNα or left non-stimulated (3.6 × 10^5^ cells, triplicates). After 6 h, the cells were harvested and total RNA was purified as described above. Total RNA was used to generate RNA sequencing libraries using the QuantSeq 3’ mRNA-seq library prep kit FWD (Lexogen). The samples were sequenced on a HiSeq 1500 device (Illumina). For data analysis, the reads were aligned to the human reference genome (Ensembl genome version 96) using STAR aligner, quantified with HTSeq, and differential expression analysis was performed using DESeq2 ^56–58^.

### PLA

The Duolink in situ red kit rabbit/mouse (Sigma-Aldrich) was used according to supplier’s instructions to analyze interactions between hnRNPM, ELAVL1, and pTBK1-Ser172 by PLA. Briefly, hnRNPM-GFP-, ELAVL1-GFP-, or GFP-expressing THP-1 cells were PMA-differentiated in CellCarrier-96 ultra imaging microplates (PerkinElmer), stimulated with AF647-labelled G_3_-YSD, fixed with 4% (w/v) paraformaldehyde in PBS (pH 6.9) at room temperature for 10 min, and permeabilized with 0.3% (v/v) Triton X-100-PBS at room temperature for 10 min. Following, the samples were washed twice with PBS and then blocked with 40 µl Duolink blocking solution at 37 °C for 60 min and incubated with the respective primary antibodies against two proteins on ice for 2 h. The primary antibodies were diluted in Duolink antibody diluent as follows: anti-GFP (1:200, mouse mAb), anti-phospho-TBK1 (Ser172) (1:100, rabbit mAb), anti-ELAVL1 (1:100, rabbit mAb). The specimens were washed with Duolink wash buffer A (4× 150 µl, 5 min each wash) and subsequently incubated with 40 µl Duolink PLA probes solution, containing secondary antibodies conjugated with PLA probes plus or minus. After 1 h at 37 °C, the samples were washed with Duolink wash buffer A (3× 150 µl, 5 min each wash) and incubated with 40 µl Duolink ligase solution at 37 °C for 30 min. Subsequently, the cells were washed with Duolink wash buffer A (3× 150 µl, 5 min each wash), incubated with 40 µl Duolink polymerase solution at 37 °C for 100 min, followed by washing with Duolink wash buffer B (3× 150 µl, 10 min each wash) and counterstaining of the nuclei with Hoechst 33342-PBS (5 µg/ml) for 10 min. After washing with PBS (3× 150 µl, 5 min each wash), the samples were stored in PBS at 4 °C and imaged on a Leica SP8 confocal microscope (HC PL APO CS2 63x/12.0 water immersion objective) by recording z stacks (0.5 µm z step size) in line sequential scan mode with zoom factor 4 and line average 4. PLA signals were quantified using Fiji. To reduce background signals, a threshold was set (default threshold: 6–255 or 10–255). Particles (5–∞ pixel units or 10–∞ pixel units) detected in individual cells were counted and the total number of particles for all z stack images was defined as the PLA score, providing a proportional measure to the number of protein interactions. Laser intensities, threshold settings, and the range of the particle size were fixed within one experimental group. Images are shown as maximum intensity projections.

### Quantitation and statistical analysis

Statistical significance was determined with GraphPad Prism 9. Statistical parameters are reported in the figure legends. Significances are indicated as follows: * (*P* < 0.05), ** (*P* < 0.01), *** (*P* < 0.001), **** (*P* ≤ 0.0001), ns: not significant.

### Data availability

The mass spectrometry proteomics data have been deposited to the ProteomeXchange Consortium via the PRIDE partner repository with the dataset identifier PXD028160 (Username, Password: hCIVjtes) ^59^. The 3’-mRNA sequencing data have been deposited to the GEO data repository with the dataset identifier GSE184273 (https://www.ncbi.nlm.nih.gov/geo/query/acc.cgi?acc=GSE184273, enter token izclacmapvezdsn into the box).

### Code availability

Requests for code should be directed to and will be fulfilled by the Lead Contact, Martin Schlee.

## ACKNOWLEDGEMENTS

We thank Saskia Schmitz, Christina Wallerath, and Laura Mlitzko for technical assistance. Preparatory experiments were performed by the Core Facility Mass Spectrometry, Institute of Biochemistry and Molecular Biology, Medical Faculty, University of Bonn with a mass spectrometer that was funded by the Deutsche Forschungsgemeinschaft (DFG, German Research Foundation) – Projektnummer 174793735. We would like to thank the Microscopy Core Facility of the Medical Faculty at the University of Bonn for providing help, services, and devices funded by the DFG – Projektnummer 388159768. We would like to acknowledge the assistance of the Flow Cytometry Core Facility at the Institute of Experimental Immunology, Medical Faculty at the University of Bonn – Projektnummer 216372545. We would like to acknowledge the assistance of the Next Generation Sequencing Core Facility at the Institute of Human Genetics, Medical Faculty at the University of Bonn.

This study was funded by the DFG under Germany’s Excellence Strategy – EXC2151 – 390873048 of which E.B., G.H., M.G., H.K., F.I.S., and M.S. are members. It was also supported by other grants of the DFG, including TRR237 (E.B., G.H., M.S., M.G., B.M.K., C.G., H.K., F.I.S., A. Pichlmair), SFB670 (E.B., G.H., M.S.), SFB704 (G.H.), Bo&MeRanG GRK 2168 (E.B., M.S.), Emmy Noether Programme 322568668 (F.I. Schmidt). and DFG SCHL1930/1-2 (M.S.). E.B. and M.S. received financial support from BONFOR (University of Bonn). This work is part of the PhD thesis of A.K. at the University of Bonn.

## Author contributions

Conceptualization, A.K., A.-M.H., and M.S.

Methodology, A.K., M.S, A. Pichlmair, and M.G.

Formal Analysis, A.K., C.U., and T.M.S.-G.

Investigation, A.K., A.-M.H., C.U., A. Piras, R.D., A.G., J.W., K.C., P.L., R.J.B, and A.K.d.R.

Resources, B.M.K., M.H.C., F.I.S., C.G., H.K., E.B., G.H, A. Pichlmair, and M.G.

Writing – Original Draft, A.K., E.B., M.S.

Writing – Review & Editing, all authors

Visualization, A.K., C.U., and T.M.S.-G.

Supervision, M.S., A. Pichlmair, and M.G.

Funding Acquisition, M.S., E.B, G.H., M.G., B.M.K., C.G., H.K., F.I.S., and A. Pichlmair

---

**Figure S1:**
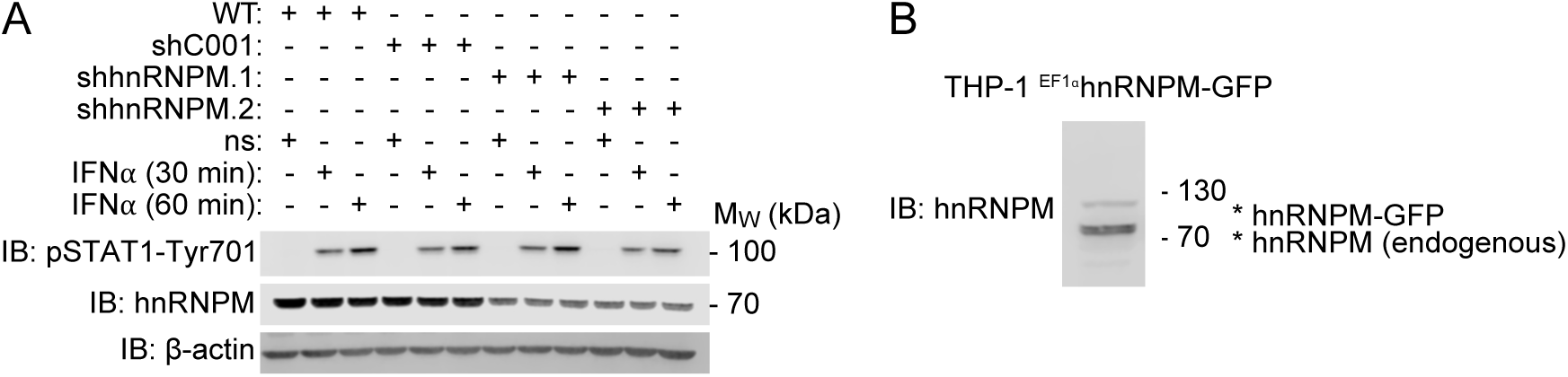
hnRNPM functions upstream of IFNAR. (A) Immunoblot analysis of pSTAT1-Tyr701 induction in THP-1 WT and cells expressing control shRNA (shC001) or hnRNPM-specific shRNAs (shhnRNPM.1, shhnRNPM.2) after stimulation with IFNα (1000 U/ml). ctrl, non-stimulated. (B) Immunoblot analysis of THP-1 cells expressing hnRNPM-GFP using an hnRNPM-specific antibody.

**Figure S2:**
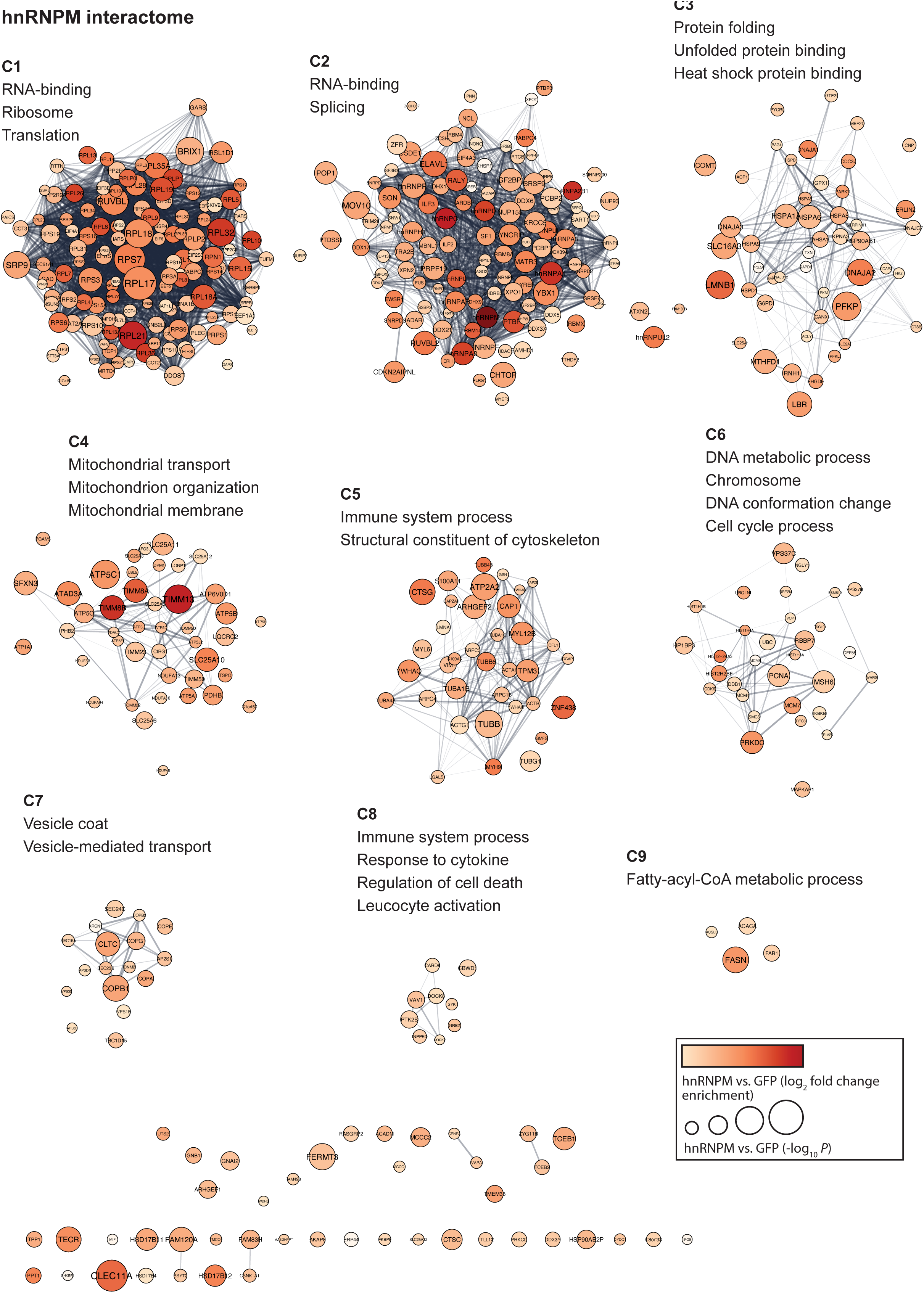
hnRNPM interacts with proteins connected to immune system processes. hnRNPM-GFP and GFP were immunoprecipitated from lysates of non-stimulated THP-1 cells. Differential interactors of hnRNPM were analyzed by STRING enrichment and annotated with GO terms enriched among hnRNPM interactors. hnRNPM interactors not annotated with these GO terms are shown at the bottom. cluster (C) 1– 9.

**Figure S3:**
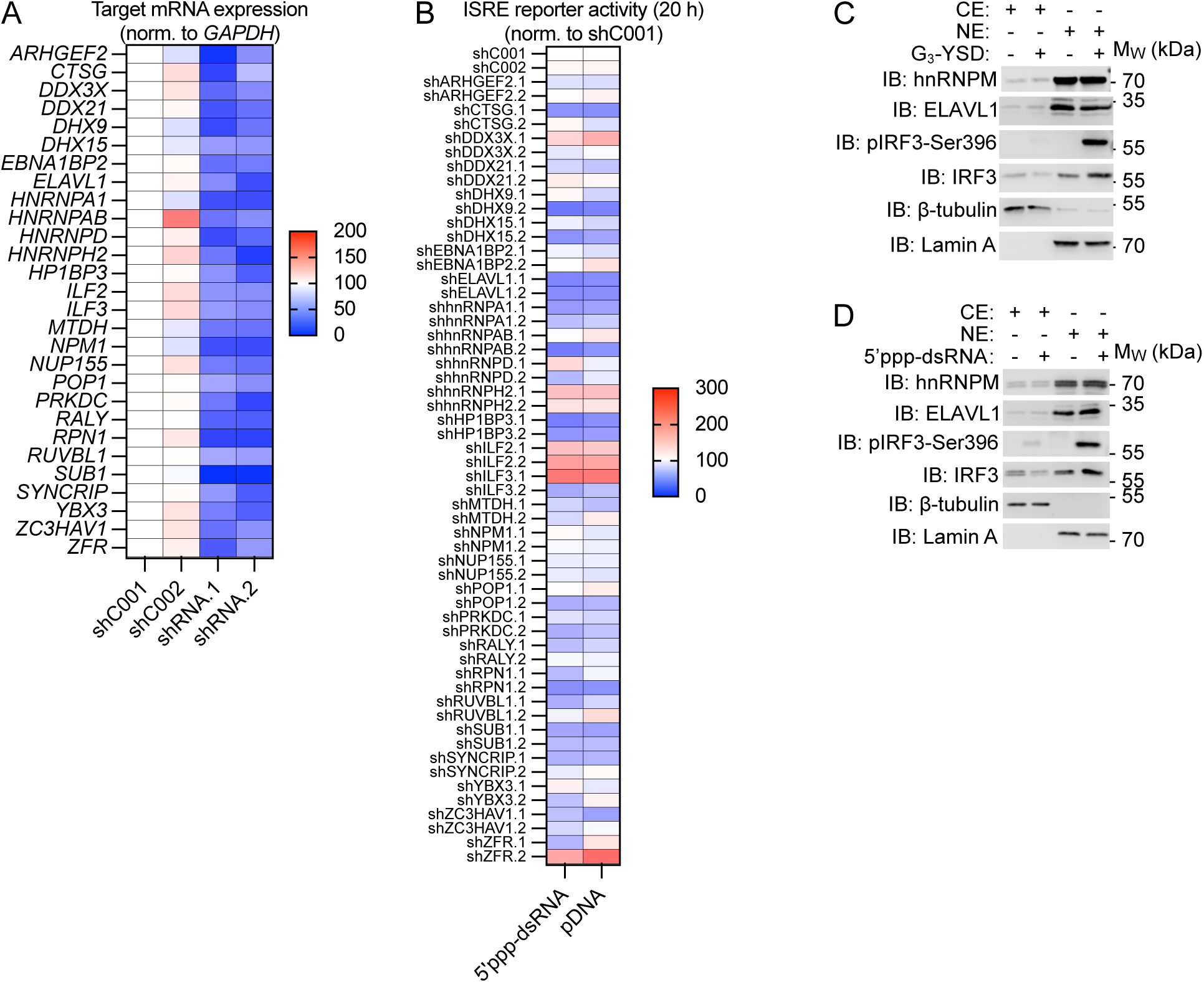
RNAi screen of hnRNPM interactors. (A) qPCR of the indicated targets (y-axis) in THP-1 cells expressing control shRNAs (shC001, shC002) or target-specific shRNAs (shRNA.1, shRNA.2). Target mRNA expression was normalized to *GAPDH* and then normalized to shC001-expressing cells. (B) ISRE reporter activation in the cells depicted in (A) 20 h after stimulation with 5’ppp-dsRNA (0.1 µg/ml) or pDNA (0.1 µg/ml). Luciferase signals were normalized to shC001-expressing cells of the respective condition. No ISRE reporter activation was detected in the non-stimulated condition (data not shown). (C–D) Nuclear extracts (NE) and cytoplasmic extracts (CE) were prepared from non-stimulated THP-1 cells or from cells stimulated with (C) G_3_-YSD (0.5 µg/ml) or (D) 5’ppp-dsRNA (0.1 µg/ml) for 3 h and analyzed by immunoblotting with the indicated antibodies. One representative experiment of two independent experiments is shown. For (A)–(B), statistical significance was determined using two-way ANOVA with Dunnett’s multiple comparisons test.

**Figure S4:**
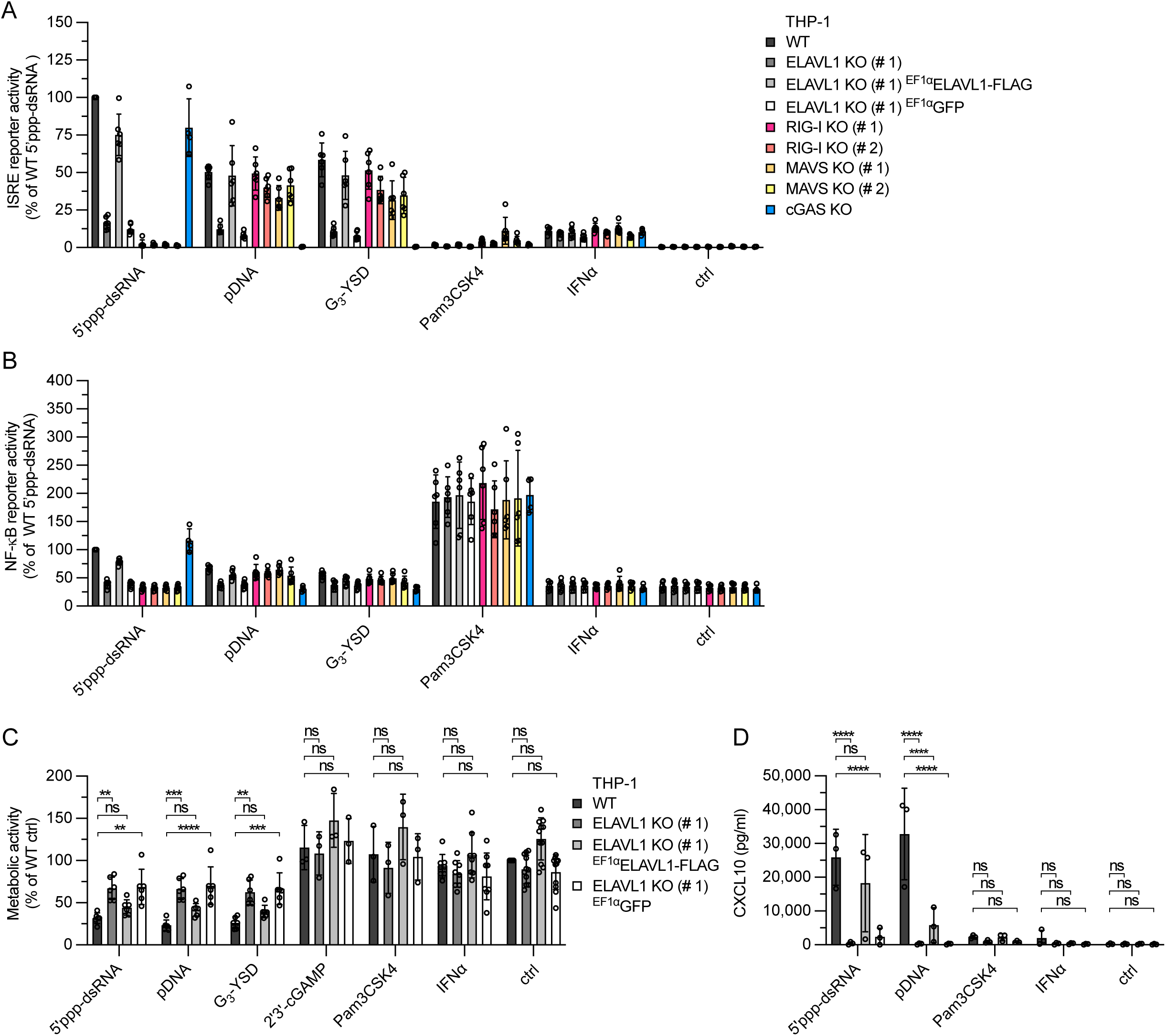
Role of ELAVL1 in cGAS and RIG-I signaling. (A) ISRE reporter activation in THP-1 WT, ELAVL1 KO (#1), ELAVL1 KO cells expressing ELAVL1-FLAG or GFP, RIG-I KO (clones #1–2), MAVS KO (clones #1–2), and cGAS KO cells 20 h after stimulation with 5’ppp-dsRNA (0.1 µg/ml), pDNA (0.1 µg/ml), G_3_-YSD (0.5 µg/ml), Pam3CSK4 (0.5 µg/ml), 2’3’-cGAMP (10 µg/ml), or IFNα (5000 U/ml) (mean ± SD). ctrl, non-stimulated. (B) NF-κB reporter activation in the cells depicted in (A) 20 h after challenge with the indicated stimuli (mean ± SD). (C) MTT assay of THP-1 WT, ELAVL1 KO (#1), and ELAVL1 KO cells expressing ELAVL1-FLAG or GFP 20 h after stimulation with 5’ppp-dsRNA (0.1 µg/ml), pDNA (0.1 µg/ml), G_3_-YSD (0.5 µg/ml), 2’3’-cGAMP (10 µg/ml), Pam3CSK4 (0.5 µg/ml), or IFNα (1000 U/ml). ctrl, non-stimulated. (D) CXCL10 ELISA with supernatants of the cells depicted in (C) collected 20 h after stimulation with 5’ppp-dsRNA (0.1 µg/ml), pDNA (0.1 µg/ml), Pam3CSK4 (0.5 µg/ml), or IFNα (1000 U/ml). ctrl, non-stimulated. For (C), (D): mean ± SD, two-way ANOVA, Dunnett’s multiple comparisons test.

**Figure S5:**
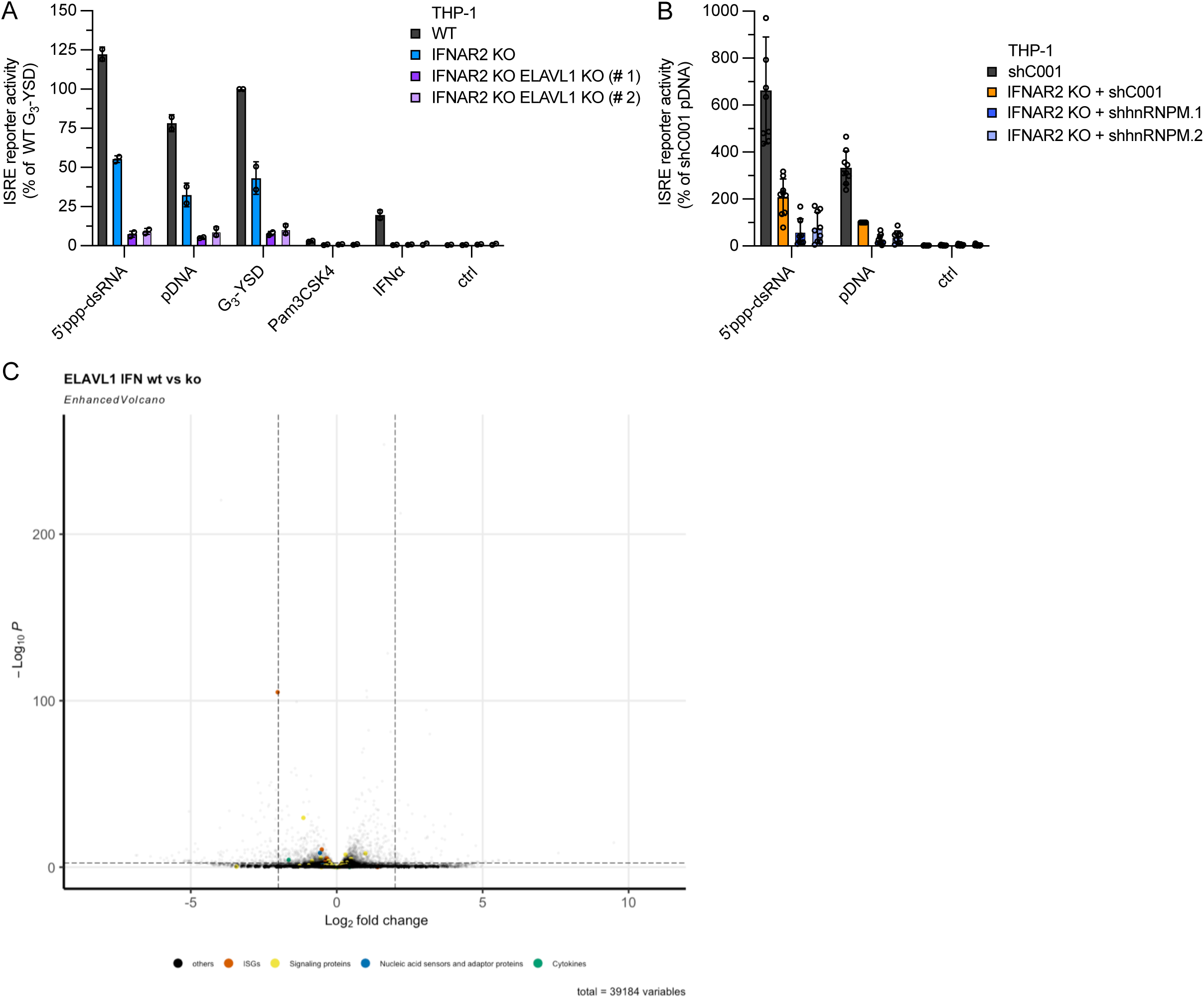
ELAVL1 regulates signal transduction downstream of cGAS and RIG-I. (A) ISRE reporter activation in THP-1 WT, IFNAR2 KO and IFNAR2/ELAVL1 double-KO cells (clones #1–2, ELAVL1 gRNA AN) 20 h after stimulation with 5’ppp-dsRNA (0.1 µg/ml), pDNA (0.1 µg/ml), G_3_-YSD (0.5 µg/ml), Pam3CSK4 (0.5 µg/ml), or IFNα (5000 U/ml) (mean ± SD). ctrl, non-stimulated. (B) ISRE reporter activation in shC001-expressing THP-1 cells and IFNAR2 KO cells expressing shC001, shhnRNPM.1, or shhnRNPM.2 20 h after stimulation with 5’ppp-dsRNA (0.1 µg/ml) or pDNA (0.1 µg/ml) (mean ± SD). ctrl, non-stimulated. (C) 3’-mRNA sequencing of total RNA from IFNα-stimulated (1000 U/ml, 6 h) THP-1 ELAVL1 KO (#1) cells and WT cells. The Volcano plot correlates the gene expression (log_2_ fold change of ELAVL1 KO vs. WT cells) with the -log_10_ adjusted *P* value (*P*_adjusted_). Significantly regulated genes were defined as *P*_adjusted_ < 0.05 and log2 fold change > 2 or log2 fold change > -2 (nucleic acid sensors and adaptor proteins (blue): cGAS, DDX58, IFIH1, MAVS, TMEM173; signaling proteins (yellow): IKBIP, TBK1, CHUK, IKBKB, IKBKE, IRF1, IRF2, IRF3, IRF4, IRF5, IRF7, IRF9, TICAM1, MYD88, TRAF1, TRAF2, TRAF3, TRAF5, TRAF6, TRAF7, TRIM25, RNF135, HMGB1, TFAM, ZCCHC3, G3BP1, NONO, IFI16, TTLL4, TTLL6, IFI16, DDX60, DHX58, IFNAR1, IFNAR2; ISGs (red): IFIT1, IFIT2, IFIT3, MX1, IL6, TNFA, IFI44L, IFI16, OASL, OAS1, OAS2, OAS3; cytokines (green): IFNB1, IFNL1, CXCL10).

